# Water as a reactant in the differential expression of proteins in cancer

**DOI:** 10.1101/2020.04.09.035022

**Authors:** Jeffrey M. Dick

## Abstract

How the abundances of proteins are shaped by tumor microenvironments, such as hypoxic conditions and higher water content compared to normal tissues, is an important question for cancer biochemistry. Compositional analysis of more than 250 datasets for differentially expressed proteins compiled from the literature reveals a higher stoichiometric hydration state in multiple cancer types compared to normal tissue; this trend is also evident in pan-cancer transcriptomic and proteomic datasets from The Cancer Genome Atlas and Human Protein Atlas. These findings support the notion of a basic physicochemical link between increased water content in tumors and the patterns of gene and protein expression in cancer. The generally increased hydration state is juxtaposed with a wide spectrum of carbon oxidation states of differentially expressed proteins, which may be associated with different gene ages, host tissue properties and metabolic features of specific cancer types.

## 1. Introduction

Although cancer is usually regarded as being driven primarily by genetic mutations [1], alterations in cancer cell metabolism and tumor microenvironment are also crucial for the growth of cancer cells. Together with extracellular acidosis due to increased glycolysis [2], changes in water and oxygen content are major chemical characteristics of cancer. Hypoxia, or less than normal physiological concentration of oxygen, in tumor microenvironments plays a major role in the biochemistry, physiology and progression of cancer [3]. Cancer tissue also has a relatively high water content [4], as consistently demonstrated by early desiccation experiments [5]. More recent developments of spectroscopic methods further substantiate the generally higher water content of cancer tissue [6,7]. These observations are consistent with the hypothesis that higher cellular hydration in carcinogenesis is a major factor that is shared with embryonic conditions [8]. Moreover, water content is a key player in other aspects of cell biology such as entry into dormancy [9]. Nonetheless, the connections between cellular water content and biomolecular abundances are not well understood.

Differences in the abundances of many proteins are a major outcome of the combination of genetic, microenvironmental and metabolic alterations in cancer. In a genocentric view of metabolism, the myriad reactions underlying changes to the proteome are catalyzed and regulated by the enzymatic products of the genome, but an adequate biochemical description should also account for the chemical compositions of the proteins themselves. From a geochemical perspective, a natural question to ask is whether the chemical compositions of differentially expressed proteins are shaped by the physicochemical conditions of tumor microenvironments.

In addition to the altered oxygenation and hydration status of tumors, the observation that most biochemical transformations involve some combination of oxidation-reduction and hydration-dehydration reactions [10,11] leads to the hypothesis that changes of oxidation and hydration state of biomolecules constitute primary biochemical variables. The sensitivity of metabolic reactions to hypoxia is well documented; for example, the reduction of metabolites under hypoxic conditions is possible by running the TCA cycle in reverse [12], and hypoxic regions in tumors accelerate the reduction of nitroxide, a redox-sensitive contrast agent used in magnetic resonance imaging [13,14]. However, hypoxia also induces the mitochondrial production of reactive oxygen species [15], so it would be an oversimplification to state that hypoxia leads to uniformly more reducing intracellular conditions. Cellular hydration state also has wide-ranging effects on cell metabolism [16], but no previous studies have systematically characterized chemical metrics of oxidation and hydration state at the proteome level in cancer.

In a short section on “Water as a Reactant”, a popular biochemistry textbook [17] describes a few types of reactions involving the release of H_2_O as a product (oxidation of glucose, condensation reactions) or its consumption as a reactant (water splitting in photosynthesis, and hydrolysis, the reverse of condensation). The polymerization of amino acids is a type of condensation reaction that is fundamental to protein synthesis, but the stoichiometry of the reactions depends only on protein length; one water is lost for each peptide bond formed between any two amino acids. A more specific metric is needed to quantify the amount of H_2_O gained or lost in the differential expression of proteins with different amino acid compositions.

Without considering detailed biosynthetic mechanisms, it is possible to use compositional metrics, which are derived from the elemental composition of proteins, to quantify the net differences in the degree of oxidation (oxidation state) and hydration (hydration state) between distinct proteins. An important theoretical consideration in deriving these metrics is that, unlike oxidation-reduction reactions, hydration-dehydration reactions do not involve the transfer of electrons, that is, they are redox-neutral. This reasoning underlies the development of a compositional metric called the stoichiometric hydration state [18]. Combined with calculations of carbon oxidation state, this makes it possible to quantify compositional differences of proteins in two dimensions that are predicted to be independent indicators of environmental oxygen and water content.

My previous analysis of proteomic data provided preliminary evidence for a higher hydration state of proteomes in colorectal and pancreatic cancer [19]. That compilation of differential expression data is expanded here to include breast, liver, lung and prostate cancer. Proteomic data are also considered for laboratory experiments of hypoxia, because of its relevance to cancer [3], and hyperosmotic stress, which has not been reported for cancer cell lines, but permits testing the sensitivity of the compositional analysis to changes in hydration state. Furthermore, I separately analyze proteomic data for both cellular and secreted proteins in hypoxia compared to normoxic controls. I also consider differential expression data for 3D culture conditions; compared to 2D or monolayer growth, the formation of cell aggregates, spheroids, or organoids in 3D culture more closely represents the tissue environment [20,21]. Finally, I combine the differential expression data with gene ages to get a picture of the evolutionary trajectories of chemical composition.

By analyzing the chemical compositions derived from proteomic datasets for particular cancer types and cell culture conditions, as well as pan-cancer transcriptomic and proteomic data, I show that hyperosmotic and 3D culture conditions in laboratory experiments induce the expression of proteins with an overall lower hydration state, whereas a higher hydration state characterizes the populations of proteins that are up-regulated in most cancer types. Therefore, the differential expression of proteins in most cancer types can be characterized theoretically as an overall biochemical reaction that consumes water as a reactant; this may be a novel biochemical manifestation of the generally elevated water content in tumors. In contrast, different cancer types show a wide range of carbon oxidation states of differentially expressed proteins, which therefore do not appear to be driven by the general condition of tumor hypoxia. Instead, the changes in oxidation state can be correlated with gene ages and might be associated with specific tissue and metabolic characteristics of different cancer types.

## 2. Results

Extensive literature searches were performed to build a database of differentially expressed proteins in five cell culture conditions versus controls and primary cancers of six organs compared to normal tissue (Fig. 1). Multiple datasets for each condition and cancer type were considered in order to compensate for inevitable technical and biological variability. In total, 301 datasets were obtained from 213 studies for cell extracts in hypoxia [22–43], secreted proteins in hypoxia [31,33,39,44–59], hyperosmotic stress represented by high salt [60–70] or high glucose [71–85], 3D vs 2D cell culture [86–104], and breast [105–121], colorectal [122–147], liver [148–176], lung [177–196], pancreatic [197–217], and prostate [218–234] cancer.

**Figure 1.**
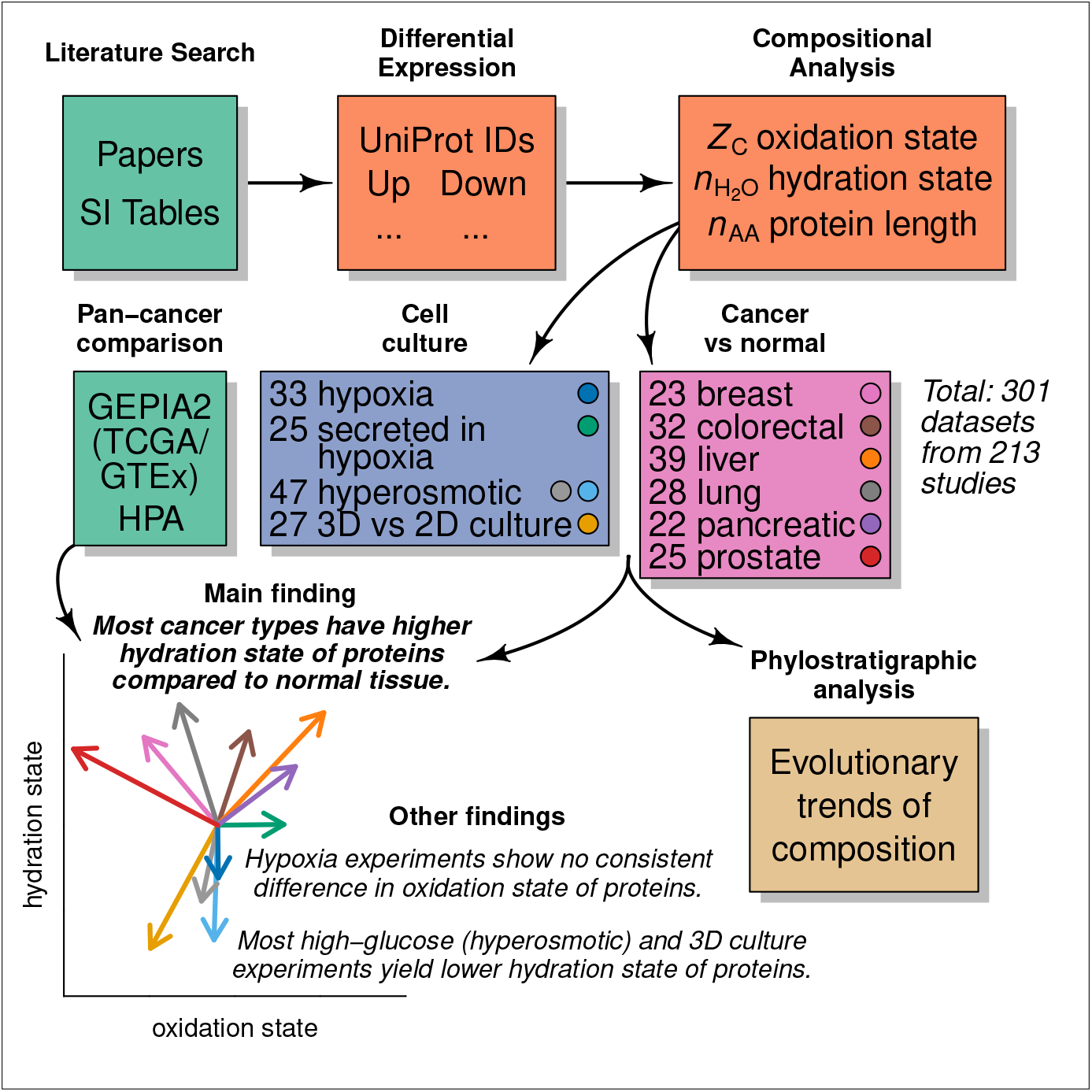
Study overview. Abbreviations: SI – Supplementary Information; GEPIA2 – Gene Expression Profiling Interactive Analysis web server; TCGA – The Cancer Genome Atlas; GTEx – Genotype-Tissue Expression project; HPA – Human Protein Atlas. The number of datasets listed for hyperosmotic conditions includes both high-salt (22) and high-glucose (25) experiments. The arrow diagram represents the mean values of differences of hydration state 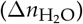 and oxidation state (Δ*Z*_C_) for all datasets in each condition (see Table 2).

The carbon oxidation state (*Z*_C_) and stoichiometric hydration state 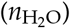 (Table 1) are compositional metrics derived from the chemical formulas of amino acids; therefore, they do not denote any particular biological mechanisms for amino acid synthesis. All of the calculations in this study are based on differences in the chemical composition of proteins as determined by their primary sequences, and do not take account of post-transcriptional modifications, like the oxidation of cysteine to make disulfide bonds, or the presence of water molecules in the hydration shell of folded proteins.

**Table 1.** Average oxidation state of carbon (*Z*_C_), number of carbon atoms (*n*_C_), and stoichiometric hydration state 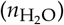 of amino acid residues computed using the rQEC derivation (see Materials and Methods and ref. [18]).

| AA | $Z_C$ | $n_C$ | $n_{H_2O}$ | AA | $Z_C$ | $n_C$ | $n_{H_2O}$ |
| --- | --- | --- | --- | --- | --- | --- | --- |
| A | 0 | 3 | 0.369 | M | -2/5 | 5 | 0.046 |
| C | 2/3 | 3 | -0.025 | N | 1 | 4 | -0.122 |
| D | 1 | 4 | -0.122 | P | -2/5 | 5 | -0.354 |
| E | 2/5 | 5 | -0.107 | Q | 2/5 | 5 | -0.107 |
| F | -4/9 | 9 | -2.568 | R | 1/3 | 6 | 0.072 |
| G | 1 | 2 | 0.478 | S | 2/3 | 3 | 0.575 |
| H | 2/3 | 6 | -1.825 | T | 0 | 4 | 0.569 |
| I | -1 | 6 | 0.660 | V | -4/5 | 5 | 0.522 |
| K | -2/3 | 6 | 0.763 | W | -2/11 | 11 | -4.087 |
| L | -1 | 6 | 0.660 | Y | -2/9 | 9 | -2.499 |

**Table 2.** Mean differences for all differential expression datasets in each condition, followed by log_10_ of *p*-value in parentheses. *p*-values less than 0.05 (log_10_ < −1.3) are shown in bold.

| Condition | $\Delta Z_C$ | $\Delta n_{H_2O}$ | $\Delta PS^a$ | $\Delta PS^b$ | $\Delta n_{AA}$ |
| --- | --- | --- | --- | --- | --- |
| *Cell culture<sup>c</sup>* |  |  |  |  |  |
| Hypoxia (cell extracts) | 0.000 (-0.0) | -0.009 (-1.6) | 0.45 (-2.5) | 0.18 (-1.2) | 7.6 (-0.1) |
| Secreted in hypoxia | 0.011 (-1.4) | 0.000 (-0.0) | 0.26 (-0.3) | 0.02 (-0.0) | -19.6 (-0.3) |
| Hyperosmotic (salt) | -0.003 (-0.5) | -0.013 (-1.5) | -0.17 (-0.7) | -0.06 (-0.1) | 16.4 (-0.4) |
| Hyperosmotic (glucose) | -0.001 (-0.1) | -0.019 (-3.6) | -0.09 (-0.2) | -0.22 (-0.4) | 11.3 (-0.3) |
| 3D / 2D | -0.011 (-2.0) | -0.021 (-4.2) | 0.14 (-0.4) | -0.08 (-0.3) | -12.6 (-0.2) |
| *Cancer<sup>c</sup>* |  |  |  |  |  |
| Breast | -0.012 (-2.7) | 0.015 (-1.7) | -1.86 (-6.4) | -0.84 (-5.9) | -71.2 (-1.8) |
| Colorectal | 0.005 (-1.1) | 0.016 (-4.3) | -0.50 (-2.0) | -0.31 (-2.7) | 36.2 (-1.0) |
| Liver | 0.018 (-6.7) | 0.019 (-5.5) | 0.37 (-1.4) | 0.54 (-4.9) | 20.5 (-0.9) |
| Lung | -0.006 (-1.1) | 0.020 (-3.4) | -1.04 (-2.8) | -0.65 (-2.7) | -44.6 (-0.9) |
| Pancreatic | 0.013 (-2.2) | 0.010 (-0.9) | 1.07 (-1.9) | 0.49 (-1.5) | 29.5 (-0.5) |
| Prostate | -0.024 (-9.4) | 0.013 (-2.0) | -1.91 (-8.9) | -1.05 (-9.5) | -25.1 (-0.4) |
| *Pan-cancer<sup>c</sup>* |  |  |  |  |  |
| TCGA / GTEx | -0.009 (-4.0) | 0.006 (-2.8) | -0.55 (-2.5) | -0.25 (-3.0) | -81.6 (-3.8) |
| HPA | -0.000 (-0.0) | 0.009 (-3.8) | 0.23 (-0.9) | -0.08 (-0.6) | 12.4 (-0.7) |
| *Secreted in hypoxia compared to cell extracts in hypoxia<sup>d</sup>* |  |  |  |  |  |
| up-regulated | 0.014 (-2.8) | -0.003 (-0.2) | 0.71 (-1.5) | 0.33 (-1.1) | 42.2 (-0.7) |
| down-regulated | 0.003 (-0.3) | -0.012 (-1.5) | 0.91 (-3.1) | 0.49 (-2.0) | 69.4 (-1.6) |

**a.** Phylostrata from ref. [240]. **b.** Phylostrata based on gene ages from ref. [241]. **c.** Differences calculated as (mean of median values for up-regulated proteins in each dataset) - (mean of median values for down-regulated proteins in each dataset), except phylostrata, where mean values for up- and down-regulated proteins in each dataset were used. The phylostrata calculations only include datasets for human proteins. $p$ -values were calculated with the paired Student's $t$ -test by using R function "t.test" [242]. **d.** Differences calculated as [mean of median values for (up- or down-)regulated proteins secreted in hypoxia] - [mean of median values for (up- or down-)regulated proteins in cell extracts in hypoxia]. $p$ -values were calculated with the unpaired Student's $t$ -test.

Carbon oxidation state for biomolecules lies between the extremes of −4 for CH_4_ and +4 for CO_2_ (see Figure 1 of ref. [235]). Because it is based on the relative electronegativities of elements, it can be calculated directly from the elemental composition of proteins [236,237]. On the other hand, a metric for hydration state depends on the stoichiometry of water in balanced chemical reactions. Since reactions that consume or release only water do not involve the transfer of electrons, a useful metric for hydration state should not be correlated with oxidation state for a collection of proteins (i.e. all those coded by the genome). Following this reasoning, the basis species glutamine–glutamic acid–cysteine–H_2_O–O_2_ were selected to write theoretical formation reactions of amino acids; the number of water molecules in these reactions (Table S1) was used as input to a residual analysis to further reduce the covariation with Z_C_ (Fig. S1), giving the residual-corrected stoichiometric hydration state listed in Table 1. This derivation, denoted “rQEC”, is briefly described in the Materials and Methods; see ref. [18] for more details and conceptual background.

### 2.1. Compositional Differences in Cell Culture Conditions and Cancer Compared to Normal Tissue

The compositional analysis of differentially expressed proteins is presented in scatterplots of median 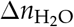 and Δ*Z*_C_, that is, the median value for all up-regulated proteins minus the median value for all down-regulated proteins in each dataset (Fig. 2 *A* and *B*). The median differences for all datasets in each condition were used to compute the 50% credible regions for highest probability density using code adapted from the “HPDregionplot” function in the R package emdbook [238], which in turn uses two-dimensional kernel density estimates calculated with “kde2d” in the R package MASS [239]. Plots with references and descriptions for all datasets are provided in Figs. S6–S16.

**Figure 2.**
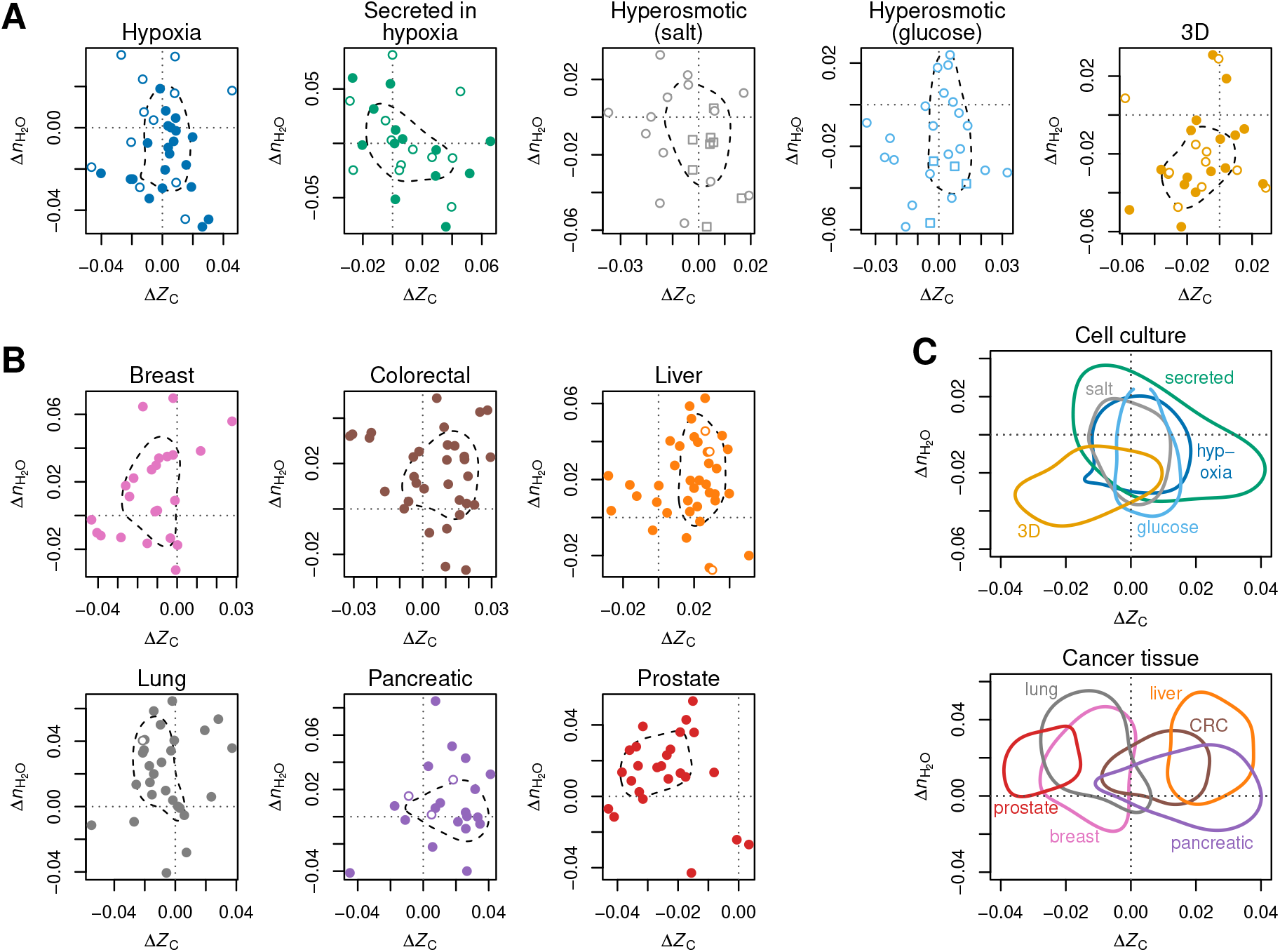
Compositional analysis of proteins identified in differential expression datasets. Median differences of stoichiometric hydration state 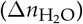 and average oxidation state of carbon (Δ*Z*_C_) in (**A**) cell culture experiments and (**B**) cancer tissues. Each point represents an individual proteomics dataset; positive value of Δ indicate a higher median value for the up-regulated proteins. For cell culture experiments, open and filled symbols represent non-cancer and cancer cells, respectively, and squares represent yeast cells. Open symbols for cancer datasets represent mouse or rat models; all others are from human subjects. Dashed lines indicate the 50% credible region for highest probability density for all datasets for each condition. (**C**) Comparison of the 50% credible regions for cell culture and cancer tissue. Abbreviation: CRC – colorectal cancer.

Several broad trends emerge from the compositional analysis of differentially expressed proteins in cell culture conditions. Differentially expressed proteins reported for cell extracts under hypoxia do not show consistent differences in oxidation state (Fig. 2*A*). However, differentially expressed proteins in many datasets for secreted proteins in hypoxia are somewhat oxidized (Δ*Z*_C_ > 0). Although the wider credible region for secreted proteins indicates a larger variability (Fig. 2*C*), the shift toward higher *Z*_C_ is statistically significant for these datasets (Table 2). Hyperosmotic stress results in the formation of proteins with predominantly lower hydration state 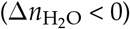; the effect is stronger for high-glucose experiments than for high-salt experiments. Lower hydration state also characterizes the majority of 3D cell culture experiments, which in addition tend to have more reduced proteins (Δ*Z*_C_ < 0). All of the 3D cell culture experiments analyzed here are for human or mouse cells, including some cancer cell lines, which are represented by filled circles in Fig. 2*A*. The experiments for cellular and secreted proteins in hypoxia include human and other mammalian cells. In contrast, the hyperosmotic stress experiments include mammalian as well as yeast cells; the latter are indicated by the squares in Fig. 2*A*. See the legends of Figs. S6–S10 for details about cell types and culture conditions.

There is a clear trend of increased hydration state of proteins 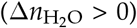 for five of the six cancer types for which multiple proteomic datasets were compiled (Fig. 2*B* and *C*). The exception is pancreatic cancer, where the datasets are distributed more evenly among positive and negative 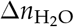. There are distinct trends in oxidation state of proteins for different cancer types: relatively oxidized proteins are up-regulated in colorectal, liver, and pancreatic cancer, whereas more reduced proteins are up-regulated in breast, lung, and prostate cancer.

The trends described above are also visible in the arrow diagram in Fig. 1. In this diagram, the lines are drawn from the origin to the mean difference of *Z*_C_ and 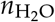 among all datasets for each cancer type and cell culture condition. The mean differences and *p*-values are listed in Table 2.

All cancer types have positive mean 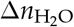, indicating greater hydration state of the up-regulated proteins, but the difference for pancreatic cancer is less statistically significant (p-value > 0.05). In contrast, hyperosmotic stress and 3D cell culture conditions, and to a lesser degree, cell extracts in hypoxia, show the up-regulation of proteins with significantly lower hydration state.

### 2.2. Elevated Hydration State and Variable Oxidation State in Pan-Cancer Datasets

Compositional analysis of large-scale pan-cancer datasets is important for assessing the differences between cancer types, and for inquiring if changes in compositional metrics for proteins are reflected in the differential expression of the genes that code for the proteins. To characterize pan-cancer transcriptomes and proteomes in terms of chemical composition, I obtained data for differential gene expression between normal tissue and cancer from GEPIA2 [245], which uses pre-compiled data files from UCSC Xena [246] that are in turn derived from the Genotype-Tissue Expression project (GTEx) [247] and The Cancer Genome Atlas (TCGA) [248]. I used data from the Human Protein Atlas (HPA) [244,249] to calculate differential protein expression as described in the Materials and Methods.

Except for prostate cancer, both pan-cancer datasets exhibit a positive 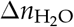 for the cancer types for which differential expression data were compiled in this study (color-coded circles in Fig. 3). Differential gene expression for all cancer types taken together corresponds to significantly more reduced proteins (Table 2, column Δ*Z*_C_), but this is not evident in the HPA proteomics datasets. In a pairwise comparison of transcriptomic and proteomic datasets for cancer types, there is very little correlation in *Z*_C_ of proteins and even less in 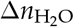 (Fig. S2). It is therefore remarkable that both the pan-cancer transcriptomic and proteomic datasets have a strong visible and statistically significant trend toward higher hydration state of the associated proteins (Table 2, column 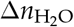. This could be an indication of an underlying genetic and biochemical tendency in cancer for increased expression of proteins with higher hydration state, despite the well-known overall weak correlation between gene and protein expression levels in cancer and other cells [250,251].

**Figure 3.**
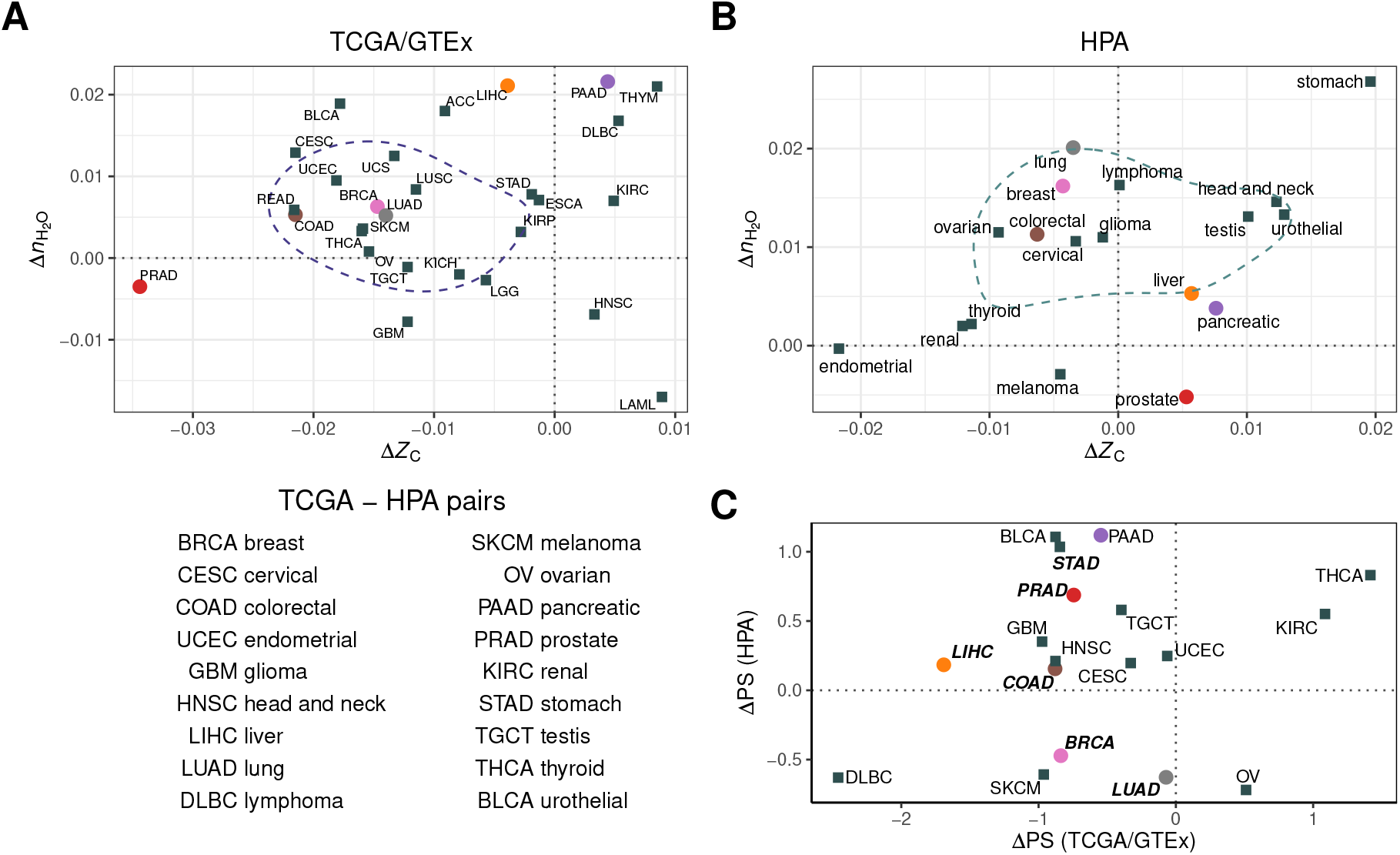
Changes in chemical composition and phylostrata for differentially regulated proteins associated with large-scale transcriptomics and protein antibody studies. (**A**) Proteins coded by differentially expressed genes between normal tissue (GTEx) and cancer (TCGA). Abbreviations for cancer types are listed in Table S2 (from ref. [243]). (**B**) Differentially expressed proteins in the Human Protein Atlas [244]. Dashed lines in panels A and B indicate the 50% credible region for highest probability density. (**C**) Mean differences of phylostrata (PS, from ref. [240]) for differentially expressed genes (TCGA/GTEx) and genes coding for differentially expressed proteins (HPA). The figure legend shows the TCGA–HPA pairings used for this plot. Color-coded circles represent cancer types with proteomic data compiled in this study (see Fig. 2) and bold italic labels indicate cancer types in the study of Trigos et al. [240]. See Figs. S17–S18 for ΔPS calculated for all TCGA and HPA datasets.

A thermodynamic mass-action hypothesis, i.e. that the nominally reducing effects of hypoxia in tumors should in general lead to greater expression of more reduced proteins, is not supported by the distribution of positive and negative Δ*Z*_C_ in the analysis of the HPA datasets. Therefore, a biological explanation is needed. It may be helpful to consider differences in the chemical composition of proteins in different subcellular locations. Membrane and extracellular proteins in yeast are relatively reduced and oxidized, respectively [237]. Similarly, stoichiogenomic analysis of the proteomes of twelve eukaryotic organisms indicates that extracellular proteins have a relatively low hydrogen content [252], which would tend to increase the average carbon oxidation state. Furthermore, up-regulated proteins that are secreted in hypoxia are more oxidized than their counterparts in cell extracts (Table 2). Given these general subcellular differences, it is interesting that tissues found by Uhlén et al. [244] to be enriched in membrane proteins (brain and kidney) host cancers with negative values of ΔZ_C_ (glioma and renal, respectively), while tissues with high levels of proteins known to be secreted (pancreas) or enriched in the transcripts of secreted proteins (liver, stomach) host cancers characterized by positive values of ΔZ_C_. These patterns seem to imply that the normal enrichment of subcellular protein classes in different tissue types could be magnified in cancer.

It is also informative to compare the compositional metrics with experimental measurements of the hydration status in different types of cancer. For instance, NMR T_1_ relaxation times are correlated with the early stages of progression of pancreatic ductal adenocarcinoma in mice, but not later stages; this is likely a consequence of increased water and protein content in the early stages [253]. This stage-specific variation of water content may help explain why the range of 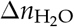 of proteins in pancreatic cancer is closer to zero, compared to other cancer types (Figs. 2*C*, 3*B*). In another study, optical measurements of gliomas in rats in the spectral range 350–1800 nm were used to infer increased water content in early stages, but decreased amounts in advanced stages, in conjunction with the formation of necrotic regions in the tumor [254]. This appears to be consistent with the small decrease in 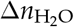 between LGG (brain lower grade glioma) and GMB (glioblastoma multiforme) in the TCGA dataset (Fig. 3*A*), but it is not possible to observe this trend in the HPA dataset, which only represents a single glioma category with positive 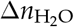 (Fig. 3*B*).

Compared to other cancer types, prostate cancer has distinct trends in the chemical composition of differentially expressed proteins. The negative 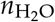 of proteins for prostate cancer in the TCGA and HPA datasets (Fig. 3) could be associated with the lower water content of prostate cancer than surrounding normal tissue, as reported in one study using near infrared spectroscopy [255]. However, for unknown reasons this trend is not apparent in the compiled proteomic datasets (Fig. 2B). Furthermore, the highly negative ΔZ_C_ of the proteins in the compiled proteomic datasets (Fig. 2B) and those coded by differentially expressed genes in prostate cancer (Fig. 3A) might somehow be related to the hypoxic characteristics of normal prostate tissue [256] and the unusual metabolic profile of prostate cancer, including de novo lipid synthesis [257] and associated impacts on cellular redox balance [258].

### 2.3. Relations between Phylostrata, Chemical Composition, and Protein Length

Several studies have linked gene expression in cancer to phylogenetically earlier genes [240,259]. The phylostratigraphic analysis used in these studies assigns ages of genes based on the latest common ancestor whose descendants have all the computationally detected homologs of that gene. To analyze the evolutionary trends of oxidation and hydration state of proteins, I used 16 phylostrata (PS) for human protein-coding genes given by Trigos et al. [240]. Using average values for proteins coded by genes in each phylostratum, Fig. 4*A* shows an initial rise in protein length, leading up to Eukaryota, which is consistent with earlier reports that median protein length is greater in eukaryotes than prokaryotes [260]. The decrease of protein length in later phylostrata is likely an artifact of BLAST-based homology searches [261].

**Figure 4.**
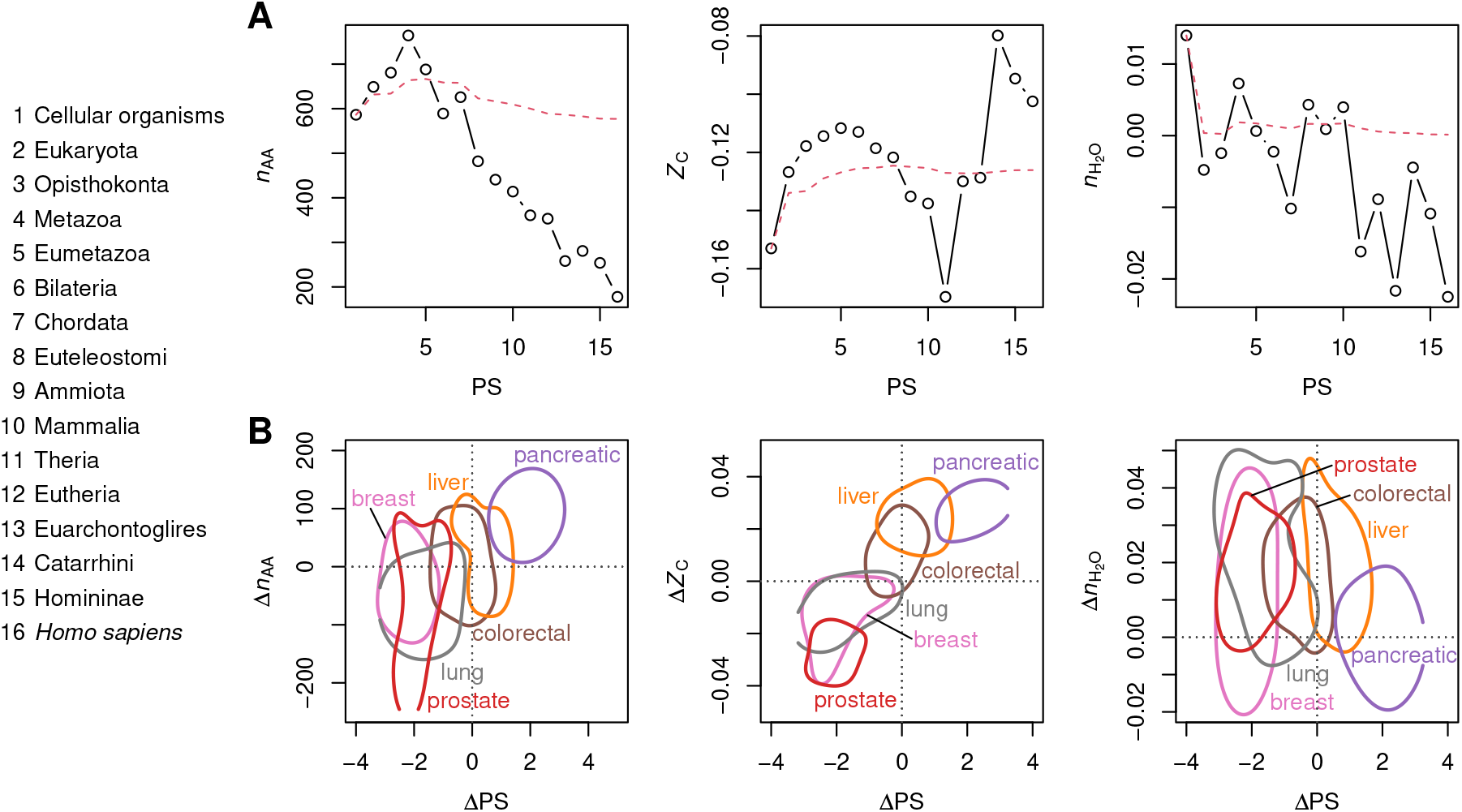
Compositional analysis of proteins using phylostrata given by Trigos et al. [240]. (**A**) Mean values of *n*_AA_, *Z*_C_, and 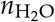 of proteins for all protein-coding genes in each phylostratum (PS). The points stand for the mean values for individual phylostrata, and the red line indicates the cumulative mean starting from PS 1. (**B**) 50% credible regions for mean differences of PS plotted against median differences of *n*_AA_, *Z*_C_, and 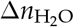 for the six cancer types for which differential expression datasets were compiled in this study. Positive values of ΔPS indicate up-regulation of proteins coded by younger genes.

Fig. 4*A* also shows distinct evolutionary patterns of oxidation state and hydration state of proteins. *Z*_C_ forms a strikingly smooth hump between PS 1 and 11 then increases rapidly to the maximum at PS 14, followed by a smaller decline to *Homo sapiens*. 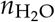 shows an overall decrease through time, but exhibits considerable local variation.

Trigos et al. [240] used RNAseq gene expression levels from TCGA for cancer and normal tissue as weights to calculate the transcriptome age index (TAI). The TAI was lower for the seven considered cancer types (LUAD, LUSC, BRCA, PRAD, LIHC, COAD, STAD; see Table S2 for definitions of abbreviations) compared to normal tissue, indicating generally higher expression of older genes. I used a different calculation, where ΔPS represents non-weighted differences between the means for up- and down-expressed genes, and obtained negative values for the same cancer types using the TCGA/GTEx data (see Fig. 3*C* for selected cancer types that are paired between TCGA and HPA and Fig. S16 for all TCGA cancer types for which differential gene expression data are available from GEPIA2, including LUSC). Therefore, in agreement with Trigos et al. [240], most cancer transcriptomes are characterized by higher expression of older genes (ΔPS < 0). However, analysis of the HPA datasets shows that many proteomes exhibit younger ages of the corresponding genes (ΔPS > 0) (Fig. 3*C*). Taken together, the negative differences for transcriptomes are more statistically significant (Table 1). The positive ΔPS for kidney renal clear cell carcinoma (KIRC) in both datasets (Fig. 3*C*), which was not analyzed by Trigos et al. [240], is consistent with the large enrichment of vertebrate genes in this cancer type [259]. The agreement with previous work suggests that ΔPS is a reasonable metric for comparing gene ages in different cancer types.

The positive association between ΔPS and protein length (Δ*n*_AA_) for differentially expressed proteins in cancer (Fig. 4*B*) implies that the corresponding genes are mostly present in Trigos PS 1 (cellular organisms) to 4 (metazoa), which are characterized by a smoothly increasing protein length (Fig. 4*A*). This agrees with previous studies wherein the differentially expressed genes in cancer were shown to consist for the most part of genes originating from the unicellular–multicellular transition [240,259]. It follows that ages inferred for cancer-related genes are not greatly affected by the artifacts that are likely present in later phylostrata assignments. To gather more evidence, I also used phylostrata corresponding to eight gene ages reported by Liebeskind et al. [241] based on consensus tables for different age-estimation algorithms. Note that the Liebeskind ages have three steps between cellular organisms and Eukaryota, providing a greater resolution in earlier evolution, and stop at Mammalia, which corresponds to Trigos PS 10. Keeping in mind the different resolutions and scales of the Trigos and Liebeskind gene ages, the two datasets show similar maxima for *Z*_C_ and protein length near Eumetazoa (or Opisthokonta, which is not one of the Trigos phylostrata), and an overall decrease of 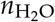 during evolution (Figs. 4*A* and S3).

A positive association between ΔPS and Δ*Z*_C_ for differentially expressed proteins in cancer would be expected based on the continually increasing *Z*_C_ for phylostrata 1–4 shown in Fig. 4*A*. Although the TCGA datasets exhibit a weak negative correlation between ΔPS and Δ*Z*_C_, the correlation is positive and somewhat stronger for the HPA datasets (Fig. S4). Similarly, a positive association between ΔPS and Δ*Z*_C_ is visible for the compiled proteomic datasets in Fig. 4*B*. It follows that differences in evolutionary ages of cancer-associated genes may help explain some of the wide variability of oxidation state of differentially expressed proteins found for different cancer types. However, the hydration state of the same proteins is largely independent of ages of the corresponding genes (Fig. 4*B*), and the magnitude of 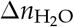 is larger in cancer (from 0.01 to 0.02; Table 2) than the cumulative difference between PS 1 and 4 (ca. 0.01; Fig. 4*A*), so something else is needed to account for the increased hydration state of proteins in cancer.

## 3. Discussion

The evolutionary trends inferred from phylostrata may help to rationalize some features of differential expression of proteins in cancer. For instance, despite the hypoxic nature of tumors, previous authors did not find significantly lower oxygen contents of proteins in glioma and stomach cancer compared to normal tissue [262,263]. Likewise, in this study a range of differences in carbon oxidation state was documented, from very negative values for prostate cancer to positive values for colorectal, pancreatic, and liver cancer. Across these cancer types, there is a close association between Δ*Z*_C_ and ΔPS (Fig. 4*B*). Unlike the proteomic data, pan-cancer transcriptomes show higher expression of generally older genes that also code for more reduced proteins (Fig. 3 *A* and *C*), so the genetic connection between the chemical compositions of proteins and tumor hypoxia appears to be shaped more by evolutionary trends than by physicochemical constraints.

It would be fruitful to compare the current results for biomolecular oxidation state with oxygen and redox measurements for tumors [13,14], but systematic measurements across tumor types may not be available. The results can also be compared with hypoxia scores computed from gene expression data [264], with the caveat that they are not physicochemical measurements. Median hypoxia scores reported for 19 tumor types [265] are not correlated with differences of *Z*_C_ or 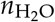 from TCGA or HPA data (Fig. S5, but note the weak positive correlation between hypoxia score and 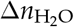 for the HPA datasets). In general, therefore, gene expression-based hypoxia scores and chemical compositions of proteins likely reflect distinct physiological processes. Nevertheless, the significantly negative Δ*Z*_C_ on average for the TCGA data (Fig. 3*A*; Table 2) raises the possibility that the chemical composition of differentially expressed proteins at the transcriptional level, but for some reason not translational level, reflects the hypoxic conditions that occur in many tumor microenvironments.

It is also somewhat surprising that hypoxia in cell culture generally does not induce the up-regulation of more reduced proteins (Table 2). The frequent downregulation of mitochondrial proteins in hypoxia [36,39] provides a possible explanation. Not only are mitochondria strongly anoxic subcellular compartments [266], but their proteins also have a relatively low *Z*_C_ compared to other subcellular fractions including the cytoplasm and nucleus [237,267], so their downregulation would tend to produce more oxidized proteins at the whole-cell level. More proteomic data for subcellular fractions are needed to better understand the overall cellular response and perhaps also what causes proteins secreted in hypoxia to be relatively oxidized (Table 2).

In marked contrast to the diverse trends of oxidation state, most cancer types are characterized by a higher stoichiometric hydration state of proteins at both the transcriptional and translational levels. These results indicate that water is consumed as a reactant when the differential expression of proteins in cancer is represented theoretically as an overall chemical reaction. This observation represents a bridge between proteomic data and experimental observations of elevated water content in tumors [6,268,269], and provides a novel line of evidence supporting the hypothesis of a primary role for elevated cellular hydration in cancer [8].

Further developments of genome-scale metabolic and macromolecular expression models [272] should be pursued to generate more precise estimates of the net water and oxygen demands for amino acid biosynthesis, uptake and incorporation into proteomes in different conditions. Nevertheless, the power of a compositional analysis of chemical formulas should not be overlooked. The quantile distributions plotted in Fig. 5 *A* and *B* provide a striking example, showing minor differences in *Z*_C_, but large positive differences of 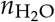 for proteins coded by up-regulated genes compared to those coded by down-regulated genes in a “common aneuploidy gene expression” dataset (CAGE) [270]. The increases of 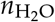 occur over a large majority of quantile points; in other words, the differences in this case exhibit nearly stochastic ordering, and therefore could arise from globally significant compositional biases [273]. Although the previous plots only show the median differences for many other datasets, it is possible to compute the quantile distributions for each dataset using the data deposited along with this paper (Section 4.5).

**Figure 5.**
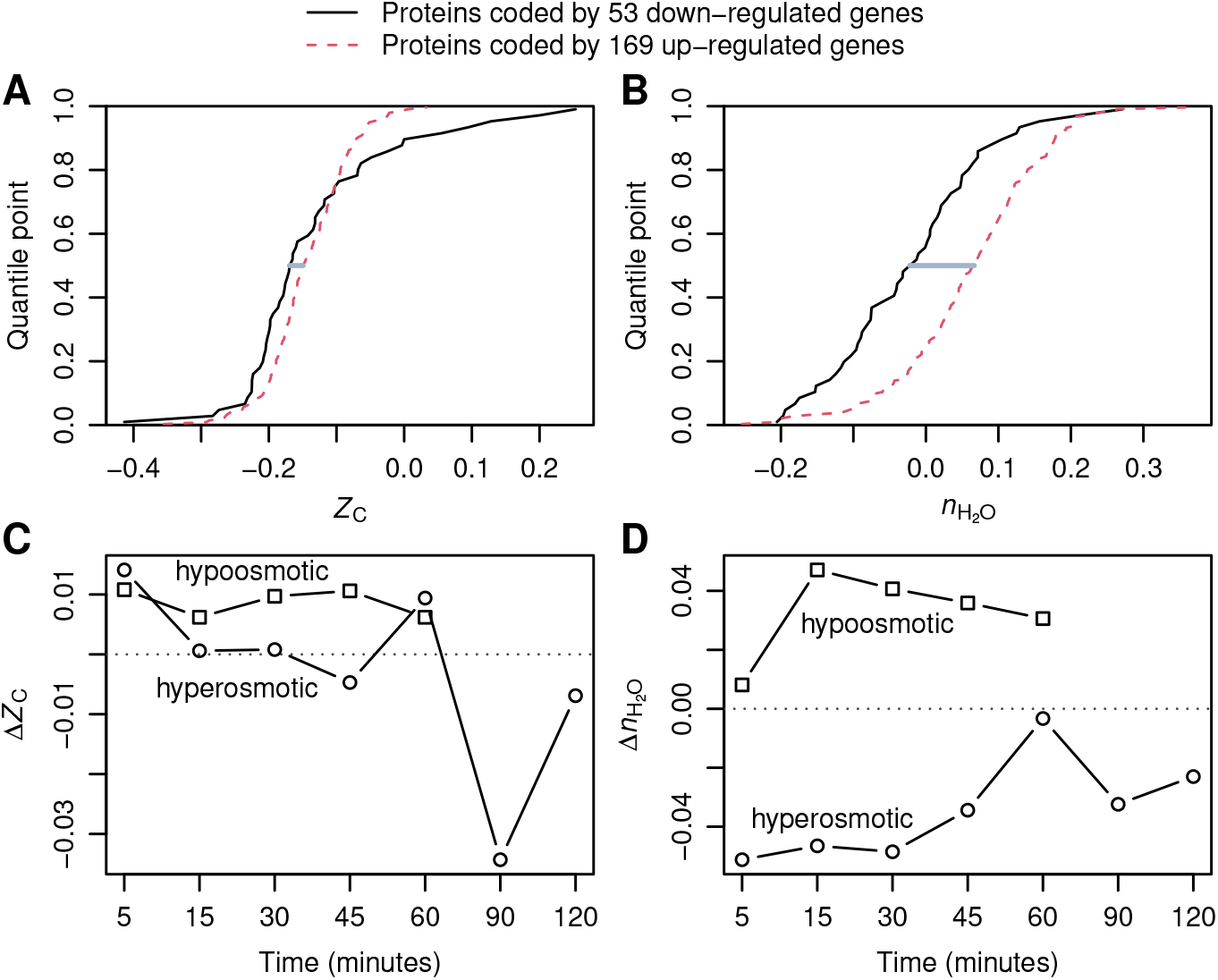
Quantile distributions of (**A**) *Z*_C_ and (**B**) 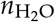 for proteins coded by differentially expressed genes in aneuploid yeast cells compared to haploid cells [270]. The median differences are indicated by the horizontal lines drawn at the 0.5 quantile point. Median differences of (**C**) Δ*Z*_C_ and (**D**) 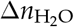 between groups of proteins coded by up- and down-regulated genes in yeast cells exposed to hyperosmotic shock (sudden shift from 0 to 1 M sorbitol) or hypoosmotic shock (growth at 1 M sorbitol followed by centrifugation to collect cells and resuspension in medium with 0 M sorbitol) [271].

Aneuploidy, or changes in chromosome number, is common in cancer cells [1], and Tsai et al. [270] found that gene expression patterns in aneuploid yeast cells are similar to those in normal yeast cells exposed to hypoosmotic (that is, more dilute, the opposite of hyperosmotic) conditions. Compositional analysis of data for gene expression in stressed yeast cells [271] indeed shows that the proteins coded by the differentially expressed genes in hypoosmotic conditions have higher 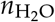, and those in hyperosmotic conditions have lower 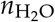 (Fig. 5 *C* and *D* and Fig. S19). The large increases in 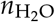 of the proteins coded by differentially expressed genes in both aneuploidy and hypoosmotic conditions suggests an possible underlying physicochemical factor: the chemical compositions of the proteins may reflect higher water activity in relatively dilute conditions. Unfortunately, gene or protein expression data for hypo- and hyperosmotic experiments have not been reported for cancer cells, so it is currently unknown whether they show a similar response.

The claim that differences of stoichiometric hydration state of proteins can be used as an indicator of physicochemical conditions of cells is strengthened by compositional analysis of proteomic data for cell culture in controlled laboratory experiments. In response to changing salinities, the interiors of most cells must be at least isosmotic with the environment to maintain a physiological water content [274]. Nevertheless, hyperosmotic conditions still exert a dehydrating effect on cell interiors, as shown by experiments with *Escherichia coli* in which the water content of cells grown in hyperosmotic NaCl solutions is substantially lowered [275]. Likewise, I found that the hydration state of differentially expressed proteins often decreases in NaCl-induced hyperosmotic stress in non-cancer eukaryotic cells (Fig. 2*C* and Table 2). A similar trend is visible in proteomics data for bacterial cells, but the effect is not very large [18]. On the other hand, experiments with eukaryotic cells in high-glucose media, which are often used to model the effects of hyperglycemia in diabetes and are also recognized for generating hyperosmotic conditions [85], show a larger average decrease in hydration state of proteins (Table 2). The present results support the hypothesis that osmotically induced dehydration provides a thermodynamic drive for the preferential expression of proteins with lower stoichiometric hydration state.

An initially unexpected finding is that the hydration state of proteins is substantially lower in 3D culture, including spheroids and aggregates, compared to traditional 2D culture in monolayers (Fig. 2*A*; see also Fig. S9). This finding might be linked with the less liquid-like state of the cytoplasm in 3D culture [276]. These results are also concordant with metagenomes of particle-sized fractions compared to free-living microbes in river and marine samples; the former, which are more likely to harbor multicellular communities, are associated with lower 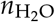 of the coded proteins [18]. Besides the strong decrease of 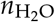, up-regulated proteins in 3D culture also tend to be more reduced (Table 2), which might reflect the attainment of hypoxic conditions in the interiors of spheroids [20,21,277].

Mesenchymal stromal cells growing in aggregates proliferate more slowly than those in monolayers under otherwise similar conditions [104]. For tumor spheroids themselves, the proliferation of cells in the core is inhibited in stress clamp experiments, which drives water out of the aggregates [278]. In monolayer cell culture, hyperosmotic conditions slow the proliferation of breast cancer cells [279] and can induce prostate cancer cells cultured at low density to enter a dormant state [280]. In contrast to lower water content, which is a feature of dormant cells [9], cell swelling in hypoosmotic conditions is a proliferative signal [281], and increased total K^+^ and water content accompanies the onset of proliferation in human blood lymphocytes exiting a quiescent state [282]. Complementary to these experimental observations, uncontrolled proliferation is identified as one of the hallmarks of cancer cells [283]. Against this background, the findings in this study of higher 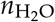 for proteins in both cancer and hypoosmotic conditions, together with lower 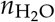 in both hyperosmotic and 3D culture conditions, could represent a new biochemical link in the association between cellular hydration levels and proliferation.

## 4. Materials and Methods

### 4.1. Proteomics Datasets

Differential protein expression data reported for any proteomics method applied to cell culture experiments and cancer compared to normal tissue were located through literature searches. Several review articles were also consulted in order to locate experimental data for breast cancer [284], lung cancer [285,286], and 3D cell culture [277]. In general, datasets were selected that have a minimum of 30 up-regulated and 30 down-regulated proteins in order to reduce random variation associated with small sample sizes, but smaller datasets (at least ca. 20 up-regulated and 20 down-regulated proteins) were included for hyperosmotic stress, secreted proteins in hypoxia, lung cancer, and prostate cancer due to limited availability of data.

Previous compilations for hypoxia and colorectal and pancreatic cancer [19] were updated in this study using more recently located datasets. Datasets related to prognosis, conditioned media, stromal samples, and adenoma were removed from the updated compilation for colorectal cancer. In addition, datasets for cellular and secreted proteins in hypoxia were considered separately, and datasets for reoxygenation after hypoxia were excluded. The previous compilation of data for hyperosmotic stress [19] was also expanded in this study, but a fish gill proteome and two transcriptomic datasets were excluded, and high-salt and high-glucose datasets were analyzed separately.

Lists of significantly differentially expressed proteins were taken directly from the original publications if possible. In cases of datasets where mass spectrometric data but not lists of differentially expressed proteins were reported, quantile normalization using function “normalize.quantiles” in the R package preprocessCore [287] was performed on the intensities or peak areas in order to obtain normalized values that were used to calculate expression ratios. Where needed, reported protein or gene identifiers were converted to UniProt IDs using the UniProt mapping tool [288]. Protein sequences downloaded from UniProt were used to generate amino acid compositions using function “read.fasta” in the R package CHNOSZ [289]. The canonical protein sequences in UniProt were used, unless isoforms were identified in the data sources. Details of additional processing steps are given with Figs. S6–S16.

### 4.2. Differential Expression from Pan-Cancer Datasets

Immunohistochemistry-based expression profiles of proteins in normal tissue and pathology samples were downloaded from the Human Protein Atlas version 19 [244,249]. Pathology and normal tissue datasets were paired based on information from the HPA web site [290]: breast cancer / breast; cervical cancer / cervix, uterine; colorectal cancer / colon; endometrial cancer / endometrium 1; glioma / cerebral cortex; head and neck cancer / salivary gland; liver cancer / liver; lung cancer / lung; lymphoma / lymph node; melanoma / skin 1; skin cancer / skin 1; ovarian cancer / ovary; pancreatic cancer / pancreas; prostate cancer / prostate; renal cancer / kidney; stomach cancer / stomach 1; testis cancer / testis; thyroid cancer / thyroid gland; urothelial cancer / urinary bladder. Antibody staining intensities were converted to a semi-quantitative scale (not detected: 0, low: 1, medium: 3, high: 5). The expression level score for each protein was calculated by averaging the score for available samples, including “not detected” but excluding unavailable (NA) observations, and, for normal tissues, observations in all available cell types. Differences in expression score between normal and cancer *≥* 2.5 or *≤* −2.5 were considered to be differentially expressed proteins.

Differential gene expression values were obtained using version 2 of the Gene Expression Profiling Interactive Analysis web server (GEPIA2) [245] with default settings (ANOVA, log_2_ fold change cutoff = 1, *q*-value cutoff = 0.01). Pairings between source datasets for cancer (TCGA) and normal tissue (GTEx), as described on the GEPIA2 website [291] are: ACC / adrenal gland; BLCA / bladder; BRCA / breast; CESC / cervix uteri; COAD / colon; DLBC / blood; ESCA / esophagus; GBM / brain; KICH / kidney; KIRC / kidney; KIRP / kidney; LAML / bone marrow; LGG / brain; LIHC / liver; LUAD / lung; LUSC / lung; OV / ovary; PAAD / pancreas; PRAD / prostate; READ / colon; SKCM/ skin; STAD / stomach; TGCT / testis; THCA / thyroid; THYM / blood; UCEC / uterus; UCS / uterus. Gene expression data for both tumor and normal tissue for HNSC are from TCGA. Differential expression data were not available on GEPIA2 for five other cancer types in TCGA (CHOL, MESO, PCPG, SARC, UVM). Ensembl Gene IDs used in HPA and GEPIA were converted to UniProt accession numbers using the UniProt mapping tool [288].

### 4.3. Compositional Metrics

Values of average oxidation state of carbon (*Z*_C_) of amino acids (Table 1) were calculated from the chemical formulas of the amino acids [236,237]. Values for *Z*_C_ of proteins were computed by combining the amino acid compositions of proteins with *Z*_C_ of amino acids and also weighting by carbon number [18]. That is, *Z*_C_ = ∑ *Z*_C,*i*_n_i_*n*_C,*i*_ / ∑ *n_i_n*_C,*i*_, where the summation is over *i* = 1..20 amino acids and *Z*_C,*i*_, *n_i_*, and *n*_C,*i*_ are the carbon oxidation state, frequency in the protein sequence, and number of carbon atoms of the *i*th amino acid, respectively.

Values of stoichiometric hydration state 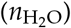 for amino acids (Table 1) were calculated using the rQEC derivation described by Dick et al. [18]. Briefly, the numbers of H_2_O in theoretical formation reactions for the 20 amino acid residues were obtained by projecting the elemental compositions of the amino acids into the basis species glutamine, glutamic acid, cysteine, H_2_O, and O_2_ (QEC basis species; see Table S1 and Fig. S1*A*). The stoichiometric hydration state was obtained by calculating the residuals of a linear model fit to 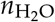 and *Z*_C_ for the amino acid residues, then subtracting a constant from the residuals to make the mean per-residue value for all human proteins equal to zero. The residual analysis ensures that there is no correlation between 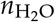 and *Z*_C_ of amino acids (Fig. S1*B*). To compute per-residue values for proteins, the values of 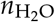 for amino acid residues from Table 1 were combined with the amino acid compositions of the proteins. That is, 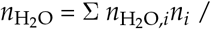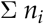, where 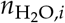 and *n_i_* are the stoichiometric hydration state and frequency of the *i*th amino acid residue, respectively. Accordingly, 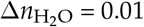 = 0.01 corresponds to a difference of approximately 3 water molecules in the theoretical formation reaction of a typical 300-residue protein.

### 4.4. Phylostrata

Phylostrata were obtained from the supporting information of Trigos et al. [240] and the “main_HUMAN.csv” file of Liebeskind et al. [241,292]. Liebeskind et al. did not give phylostrata numbers, so phylostrata 1–8 were assigned here based on the names in the “modeAge” column of the source file (see Fig. S3). The Ensembl gene identifiers in the Trigos dataset were converted to UniProt accession numbers [288]; in the case of duplicate UniProt accession numbers, the first matching phylostratum was used.

### 4.5. Data deposition

The compiled differential expression data are provided in the canprot R package, version 1.0.0 (https://cran.r-project.org/package=canprot). Other data used for this paper are in the “canH2O”, “aneuploidy”, and “yeast_stress” directories of the JMDplots package, version 1.2.2 (https://github.com/jedick/JMDplots). The specified versions of both of these packages are also deposited on Zenodo [293,294]. Figs. S6–S16 are derived from the vignettes in the canprot package, and the code for the remaining figures and Tables 2 and S1 is in the JMDplots package; the “canH2O” vignette in this package has the function calls used to make the figures and tables.

## Supplementary Materials

Figures S1–S19, Tables S1–S2.

## Acknowledgments

I am grateful to Alex Greenhough, Youngsoo Kim, Ming-Chih Lai, and Gordana Vunjak-Novakovic for providing data files. The results shown here are in part based upon data generated by the TCGA Research Network (https://www.cancer.gov/tcga) and the Human Protein Atlas (https://www.proteinatlas.org).

