## Supplementary material for "Water as a reactant in the differential expression of proteins in cancer": SI Appendix

**Jeffrey M. Dick**

****

#### This PDF file includes:

Figs. S1 to S19
Tables S1 to S2
References for SI reference citations

**Table S1. Stoichiometric matrix for amino acid residues with the QEC basis species (glutamine – glutamic acid – cysteine – $H_2O$ – $O_2$).**

| | $C_5H_{10}N_2O_3$ | $C_5H_9NO_4$ | $C_3H_7NO_2S$ | $H_2O$ | $O_2$ |
|---|---|---|---|---|---|
| alanine (A) | 0.4 | 0.2 | 0 | -0.4 | -0.3 |
| cysteine (C) | 0 | 0 | 1 | -1 | 0 |
| aspartic acid (D) | 0.2 | 0.6 | 0 | -1.2 | 0.6 |
| glutamic acid (E) | 0 | 1 | 0 | -1 | 0 |
| phenylalanine (F) | -0.8 | 2.6 | 0 | -3.2 | -1.9 |
| glycine (G) | 0.6 | -0.2 | 0 | -0.6 | 0.3 |
| histidine (H) | 1.8 | -0.6 | 0 | -2.8 | 0.4 |
| isoleucine (I) | -0.2 | 1.4 | 0 | 0.2 | -2.1 |
| lysine (K) | 0.8 | 0.4 | 0 | 0.2 | -1.6 |
| leucine (L) | -0.2 | 1.4 | 0 | 0.2 | -2.1 |
| methionine (M) | -0.4 | 0.8 | 1 | -0.6 | -1.2 |
| asparagine (N) | 1.2 | -0.4 | 0 | -1.2 | 0.6 |
| proline (P) | 0 | 1 | 0 | -1 | -1 |
| glutamine (Q) | 1 | 0 | 0 | -1 | 0 |
| arginine (R) | 2.8 | -1.6 | 0 | -0.8 | -0.1 |
| serine (S) | 0.4 | 0.2 | 0 | -0.4 | 0.2 |
| threonine (T) | 0.2 | 0.6 | 0 | -0.2 | -0.4 |
| valine (V) | 0 | 1 | 0 | 0 | -1.5 |
| tryptophan (W) | -0.2 | 2.4 | 0 | -4.8 | -1.6 |
| tyrosine (Y) | -0.8 | 2.6 | 0 | -3.2 | -1.4 |

**Table S2. Abbreviations for 33 cancer types in the TCGA PanCancer Atlas (from https://gdc.cancer.gov/resources-tcga-users/tcga-code-tables/tcga-study-abbreviations, accessed on 2020-01-30).**

| Abbreviation | Study Name | Abbreviation | Study Name |
|---|---|---|---|
| ACC | Adrenocortical carcinoma | OV | Ovarian serous cystadenocarcinoma |
| BLCA | Bladder Urothelial Carcinoma | PAAD | Pancreatic adenocarcinoma |
| BRCA | Breast invasive carcinoma | PRAD | Prostate adenocarcinoma |
| CESC | Cervical squamous cell carcinoma and endocervical adenocarcinoma | READ | Rectum adenocarcinoma |
| COAD | Colon adenocarcinoma | SKCM | Skin Cutaneous Melanoma |
| DLBC | Lymphoid Neoplasm Diffuse Large B-cell Lymphoma | STAD | Stomach adenocarcinoma |
| ESCA | Esophageal carcinoma | TGCT | Testicular Germ Cell Tumors |
| GBM | Glioblastoma multiforme | THCA | Thyroid carcinoma |
| HNSC | Head and Neck squamous cell carcinoma | THYM | Thymoma |
| KICH | Kidney Chromophobe | UCEC | Uterine Corpus Endometrial Carcinoma |
| KIRC | Kidney renal clear cell carcinoma | UCS | Uterine Carcinosarcoma |
| KIRP | Kidney renal papillary cell carcinoma | *Differential gene expression not available in GEPIA2* | |
| LAML | Acute Myeloid Leukemia | CHOL | Cholangiocarcinoma |
| LGG | Brain Lower Grade Glioma | MESO | Mesothelioma |
| LIHC | Liver hepatocellular carcinoma | PCPG | Pheochromocytoma and Paraganglioma |
| LUAD | Lung adenocarcinoma | SARC | Sarcoma |
| LUSC | Lung squamous cell carcinoma | UVM | Uveal Melanoma |

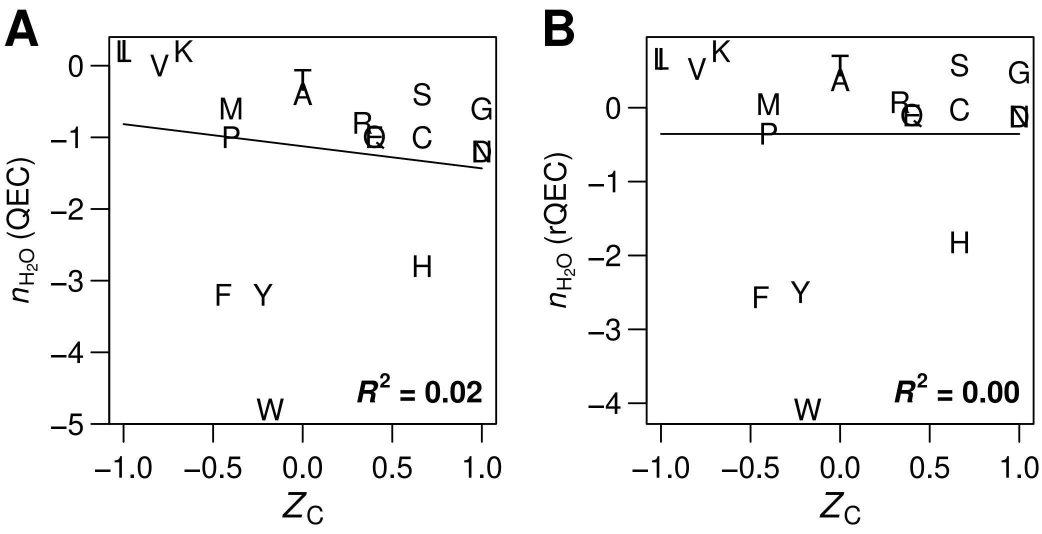


**Fig. S1.** Plots of compositional metrics for amino acid residues. Linear fits and $R^2$ values are shown. (**A**) Number of water molecules in reactions from the QEC basis species (Table S1) vs. carbon oxidation state. (**B**) Residuals from the fit in panel A, adjusted by subtracting a constant (0.355) so that the mean stoichiometric hydration state for all human proteins equals 0. The values of $n_{H_2O}$ in this plot are listed in Table 1 of the main text.

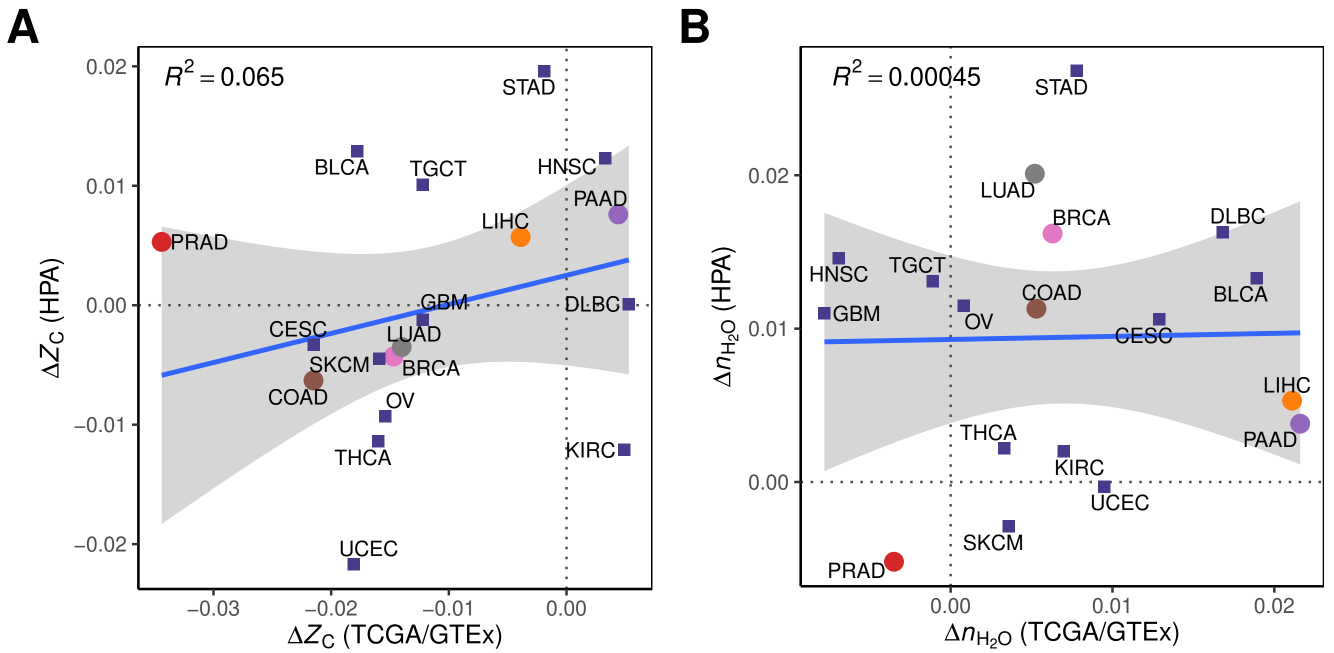


**Fig. S2.** Scatterplots of compositional metrics for differentially expressed proteins (HPA) and genes (TCGA/GTEx): (**A**) $\Delta Z_C$; (**B**) $\Delta n_{H_2O}$. Linear regressions and $R^2$ values are shown.

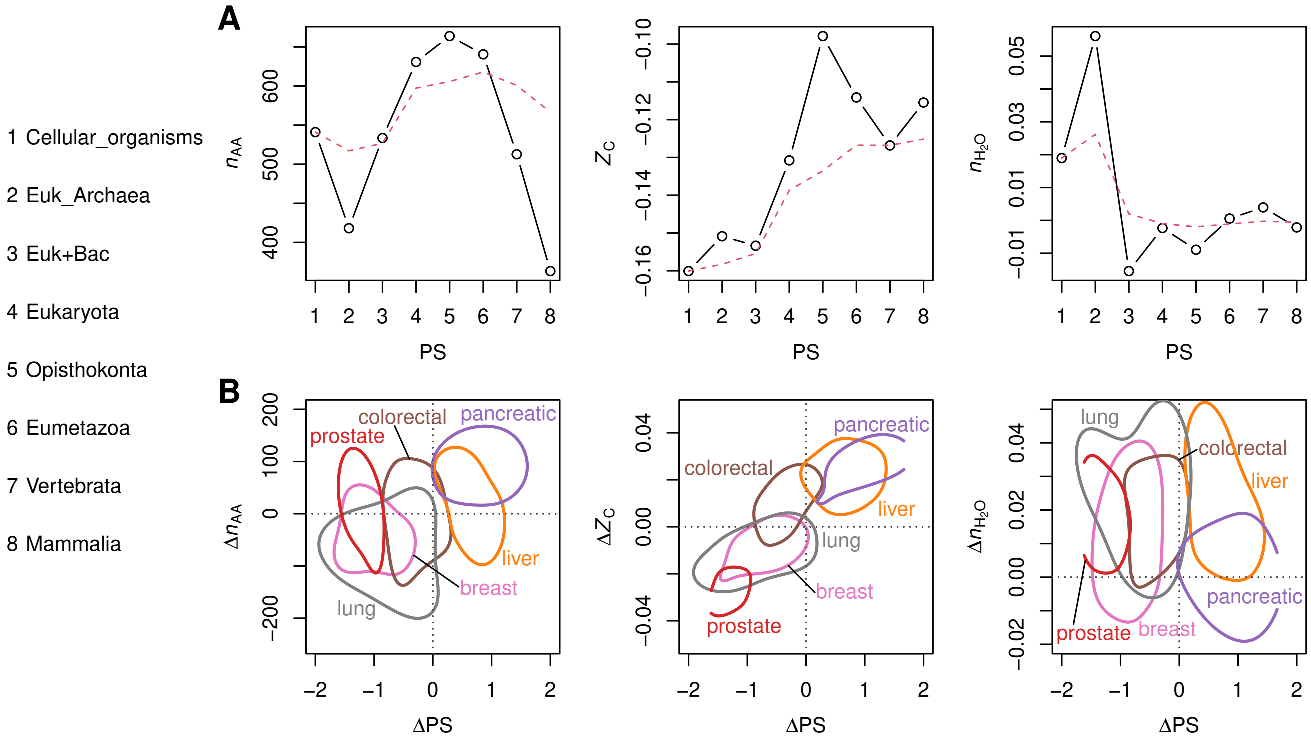


**Fig. S3.** Compositional analysis of phylostrata representing gene ages from Liebeskind et al. (1) for comparison with Fig. 4 in the main text. (**A**) Mean values of $n_{AA}$, $Z_C$, and $n_{H_2O}$ of proteins for all protein-coding genes in each phylostratum (PS). The points stand for the mean values in different phylostrata, and the red line indicates the cumulative mean starting from PS 1. (**B**) Mean differences of PS compared with median differences of $n_{AA}$, $Z_C$, and $n_{H_2O}$ for cancer datasets. The plots show the 50% credible region for differential expression datasets compiled in this study for six types of cancer.

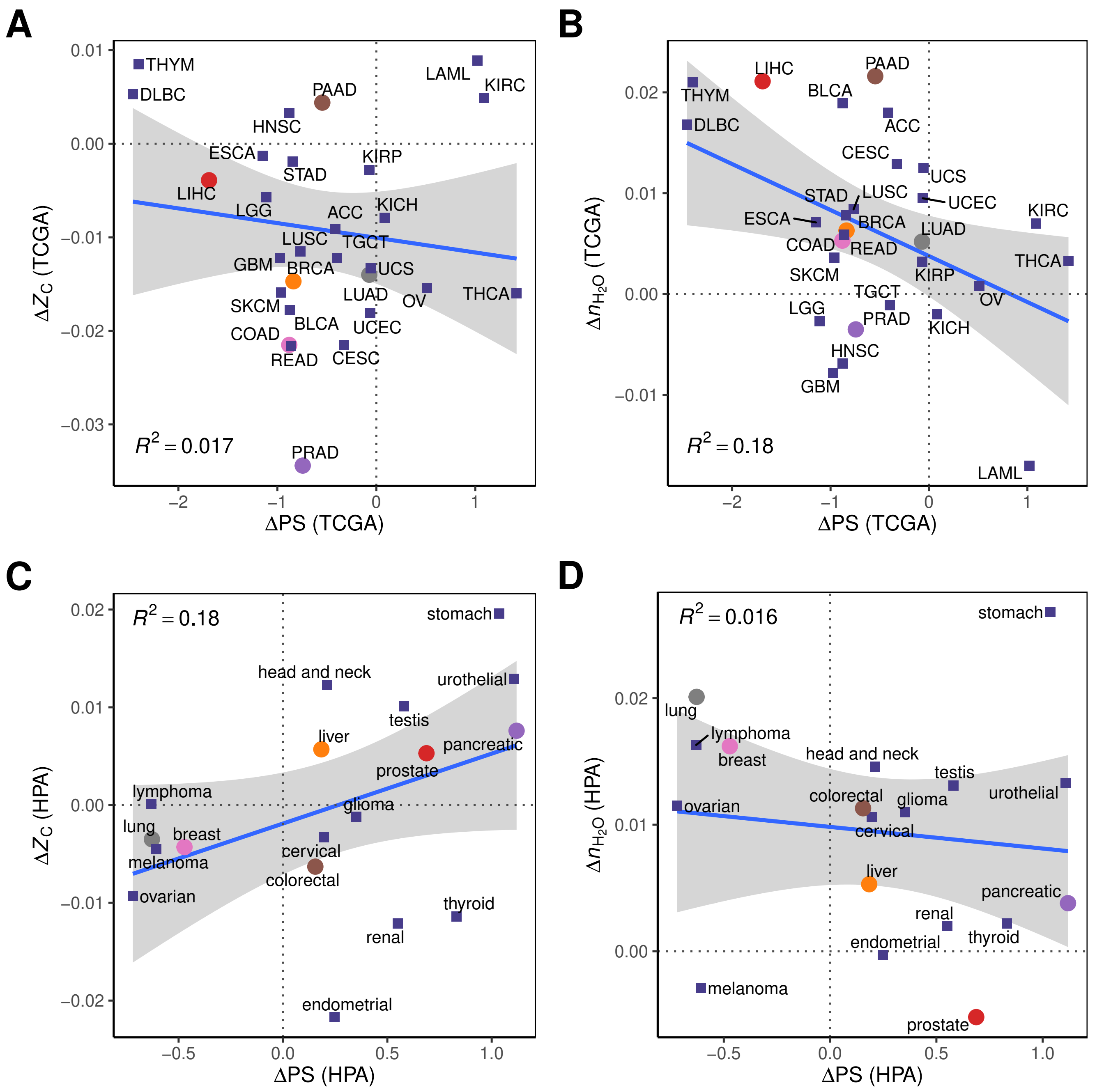


**Fig. S4.** Scatterplots for (**A**,**C**) $\Delta Z_C$ and (**B**,**D**) $\Delta n_{H_2O}$ vs ΔPS (mean phylostrata difference) for (**A**,**B**) TCGA/GTEx and (**C**,**D**) HPA datasets.

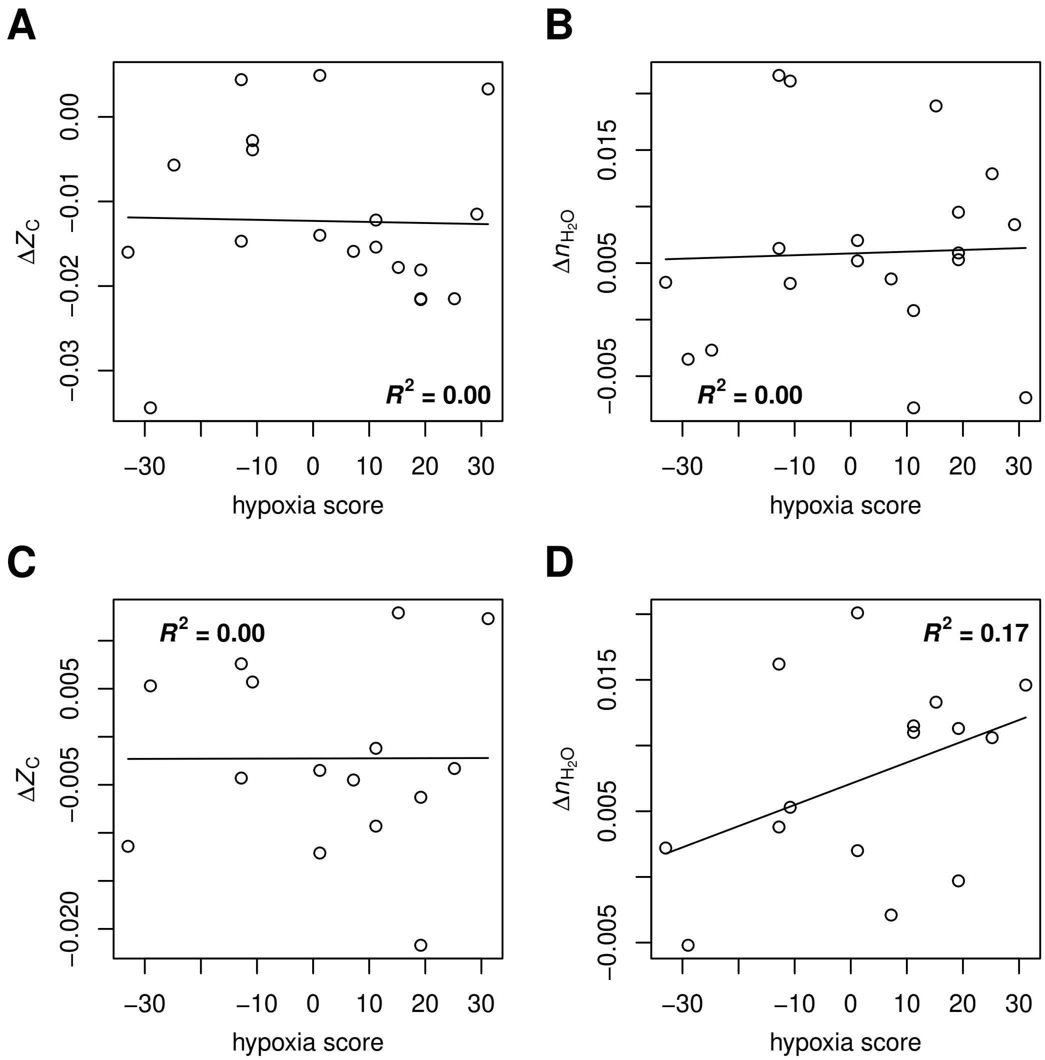


**Fig. S5.** Scatterplots of median hypoxia scores for different cancer types (2) against compositional metrics for proteins: (**A**) $Z_C$ from TCGA/GTEx datasets; (**B**) $n_{H_2O}$ from TCGA/GTEx datasets; (**C**) $Z_C$ from HPA datasets; (**D**) $n_{H_2O}$ from HPA datasets.

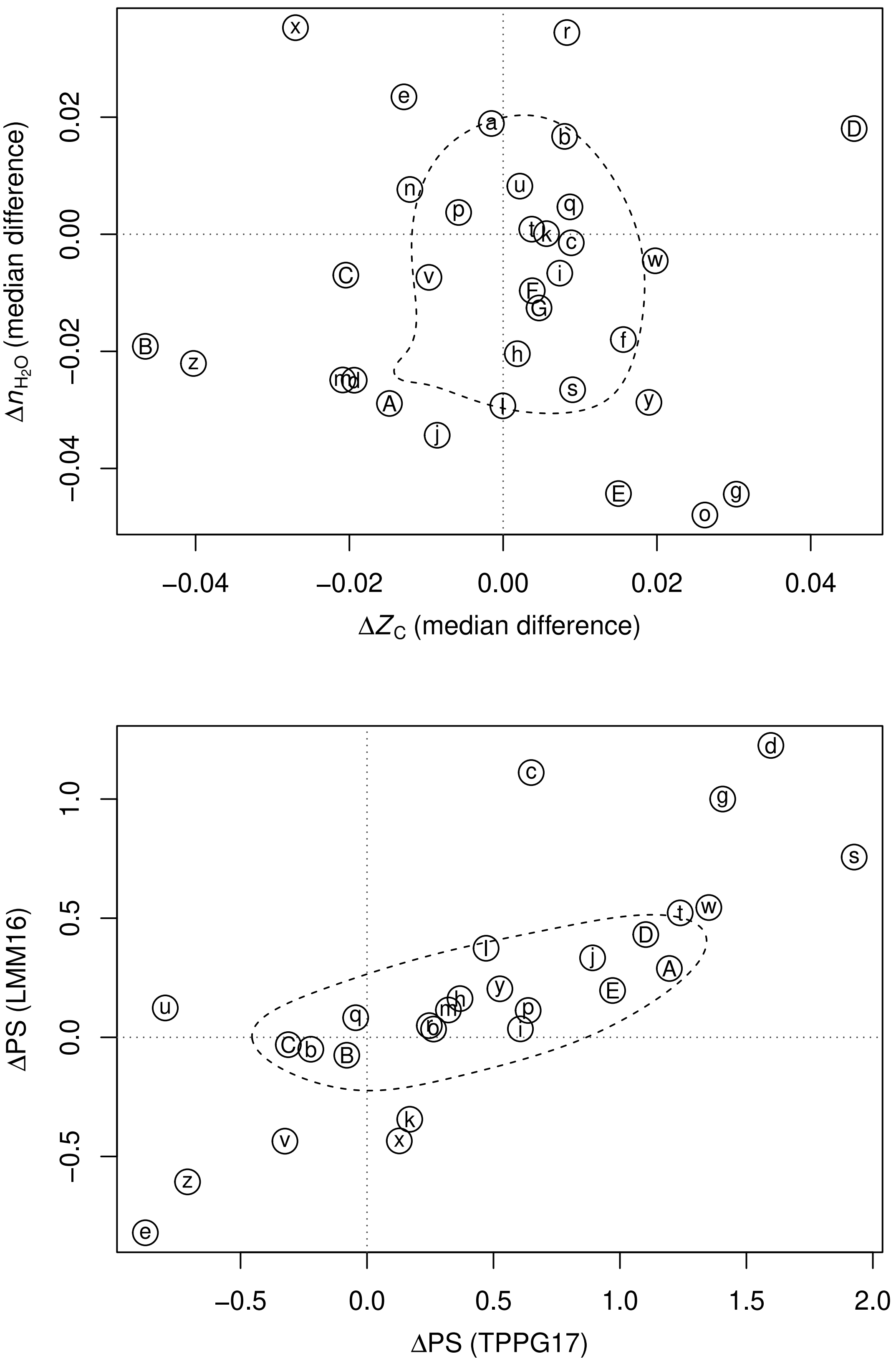


**Fig. S6.** Compositional analysis and phylostrata for cellular extracts in hypoxia.

### Legend for Fig. S6

Abbreviations: Hx48 – hypoxia 48 h; Hx72 – hypoxia 72 h; -S – supernatant fraction; -P – pellet fraction; DMSO – dimethyl sulfoxide; NO.sul – nitric oxide-donating sulindac; sul – sulindac; 4Gy – radiation; POS – primay origin canine osteosarcoma (OS) cells; HMPOS – metastatic origin canine OS cells.

| Set | Reference | Description | Down | Up |
|---|---|---|---|---|
| a | SBB+06 | mouse malignant melanoma plasma membrane | 574 | 289 |
| b | FWH+13 | THP-1 macrophages | 56 | 40 |
| c | RHD+13 | A431 epithelial carcinoma cells Hx48 | 211 | 92 |
| d | RHD+13 | A431 epithelial carcinoma cells Hx72 | 88 | 67 |
| e | VTMF13 | SH-SY5Y neuroblastoma cells | 141 | 64 |
| f | DCH+14 | mouse 4T1 cells | 71 | 60 |
| g | DYL+14 | A431 epithelial carcinoma cells Hx48-S | 65 | 34 |
| h | DYL+14 | A431 epithelial carcinoma cells Hx72-S | 137 | 61 |
| i | DYL+14 | A431 epithelial carcinoma cells Hx48-P | 74 | 44 |
| j | DYL+14 | A431 epithelial carcinoma cells Hx72-P | 67 | 53 |
| k | BSA+15 | HeLa cervical cancer cells | 53 | 72 |
| l | HWA+16 | U87MG and 786-O cancer cells | 137 | 164 |
| m | LCS16 | HCT116 colon cancer translation | 469 | 1024 |
| n | CGH+17 | mouse cardiac fibroblasts whole | 38 | 48 |
| o | ZXS+17 | U87 and U251 glioblastoma cells | 143 | 122 |
| p | CLY+18 | HCT116 colon cancer cells proteome | 108 | 127 |
| q | GBH+18 | SW620 colorectal cancer cells | 237 | 67 |
| r | LKK+18 | hUCB mesehnchymal stem cells | 44 | 62 |
| s | WTG+18 | adipose-derived mesehnchymal stem cells | 163 | 57 |
| t | CSK+19 | HeLa cervical cancer cells | 36 | 45 |
| u | GPT+19 | MIAPaCa-2 pancreatic cancer cells pulse/trace Light serum replete | 62 | 180 |
| v | GPT+19 | MIAPaCa-2 pancreatic cancer cells pulse/trace Heavy serum replete | 196 | 57 |
| w | KAN+19 | cancer-associated fibroblasts proteome | 221 | 124 |
| x | LLL+19 | human periodontal ligament cells | 67 | 153 |
| y | BCMS20 | MCF-7 breast cancer cells | 153 | 161 |
| z | RVN+20 | PC-3 prostate cancer cells in DMSO | 116 | 86 |
| A | RVN+20 | PC-3 prostate cancer cells in NO.sul | 28 | 33 |
| B | RVN+20 | PC-3 prostate cancer cells in sul | 17 | 54 |
| C | RVN+20 | PC-3 prostate cancer cells in DMSO.4Gy | 24 | 38 |
| D | RVN+20 | PC-3 prostate cancer cells in NO.sul.4Gy | 21 | 52 |
| E | RVN+20 | PC-3 prostate cancer cells in sul.4Gy | 34 | 25 |
| F | SPJ+20 | canine POS cells | 43 | 102 |
| G | SPJ+20 | canine HMPOS cells | 62 | 69 |

**a**. Supplemental Table 1 of (3), filtered to include proteins with expression ratio ≥ 1.7 or ≤ 0.58. **b**. Supplemental Table 2A of (4) (control virus cells). **c**. **d**. Supplemental Table S1 of (5), filtered to include proteins with iTRAQ ratios < 0.83 or > 1.2 and *p*-value < 0.05. **e**. Supporting Information table of (6), filtered to include proteins with a normalized expression ratio of > 1.2 or < 0.83. **f**. Supporting Information Table S1 (7). **g**. **h**. **i**. **j**. Supplemental Table S1 of (8), filtered to include proteins with *p*-value < 0.05 (-S: supernatant fraction; -P: pellet fraction). **k**. Supplementary Table S1 of (9). **l**. Supplemental Information Table S1 of (10), filtered to include proteins with a fold change of < 0.5 or > 1 and that were detected in only hypoxic or only normoxic conditions. **m**. Gene names from (11) (data files provided by Ming-Chih Lai). **n**. Extracted from Table S2E (whole cell lysate) of (12), keeping proteins with FDR < 0.05. **o**. Supplemental Table S4 of (13). **p**. Supplementary Tables S6-S7 (proteome) of (14). **q**. List of up- and down-regulated proteins from (15) (provided by Alex Greenhough), filtered to include proteins with average fold change ≥ 1.5 or ≤ 2/3. **r**. Gene names from Figure 1F of (16). **s**. Supplemental file 1 of (17) (provided by Gordana Vunjak-Novakovic), filtered to include proteins with Normalized Ratio [Hypoxia MSC/Control MSC] ≥ 1.5 or ≤ 2/3 and *p*-value < 0.05. **t**. Gene names from Supplementary file S1 of (18) for the two replicates labelled as "input_Log2ratioHL_firstIP" and "input_Log2ratioLH_secondIP" (soluble extracts before immunoprecipitation), filtered to include proteins where Log2ratio is > 0.2 or < -0.2 for both replicates. **u**. **v**. Supplementary Data 1 of (19), filtered to include proteins with *p*-value < 0.05 and fold-change > 2 or < 0.5 **w**. Proteins identified as up- or down-regulated > 1 SD in Data File S1 of (20) (sheet "Proteome"). **x**.

Additional file 1: Table S1 of (21). **y**. Gene names from Supplementary Information Tables S6 and S7 of (22). **z**. **A**. **B**. **C**. **D**. **E**. Supplementary Table 1A of (23), filtered to include proteins with median fold change between normoxic and hypoxic conditions in any individual treatment > 1.5 or < 2/3. **F**. **G**. Supplementary Tables S3b (sheet "HPNP") and S3c (sheet "HHNH") of (24).

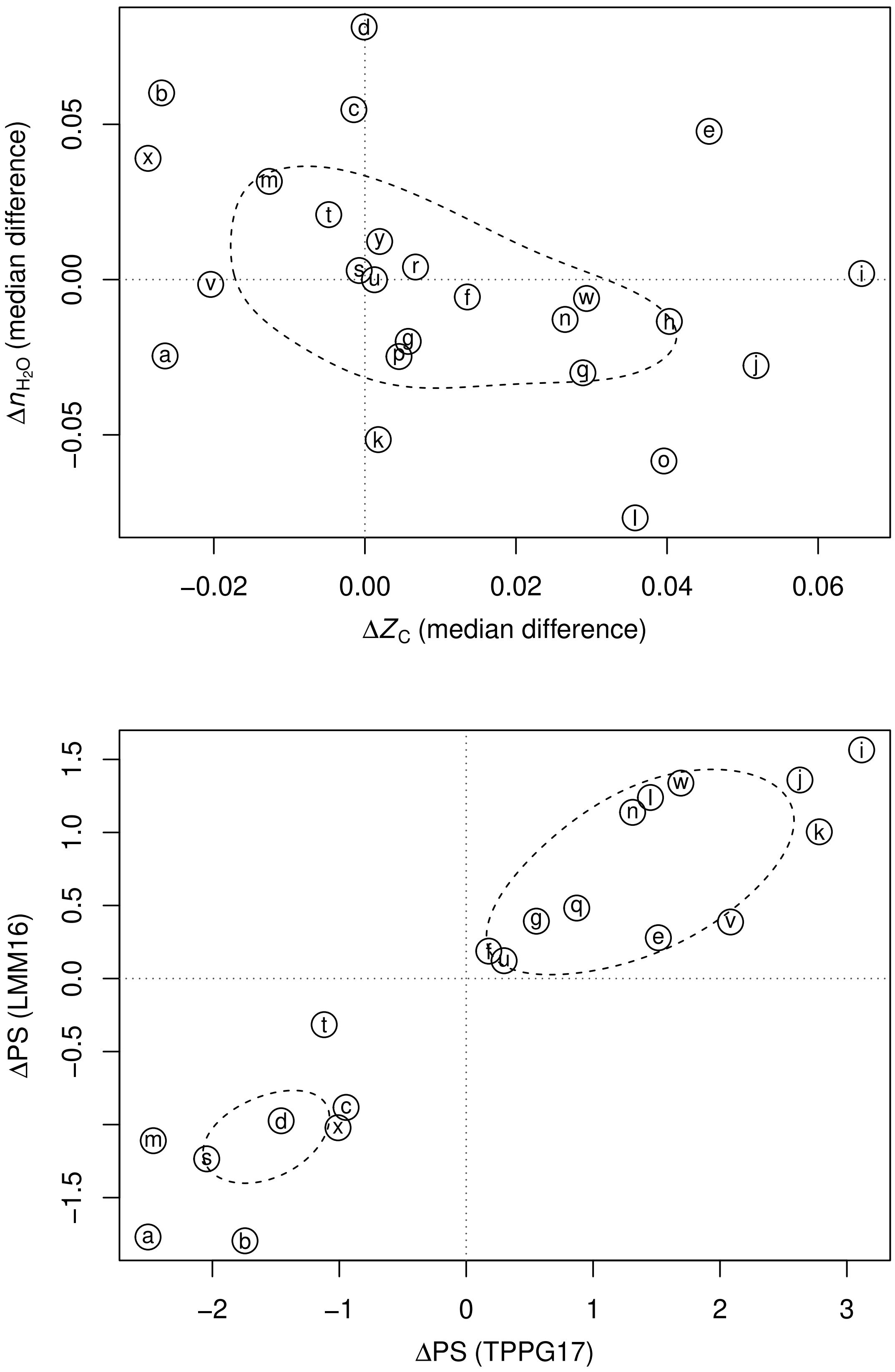


**Fig. S7.** Compositional analysis and phylostrata for secreted proteins in hypoxia.

### Legend for Fig. S7

Abbreviations: pMSC – placental mesenchymal stem cells.

| Set | Reference | Description | Down | Up |
|---|---|---|---|---|
| a | BRA+10 | placental secretome | 41 | 22 |
| b | PTD+10 | A431 squamous carcinoma cells Hx48 | 38 | 78 |
| c | PTD+10 | A431 squamous carcinoma cells Hx72 | 43 | 66 |
| d | JVC+12 | endothelial cell-derived exosomes | 64 | 45 |
| e | SKA+13 | cytotrophoblast-derived exosomes | 38 | 25 |
| f | SRS+13a | pMSC 3 / 1 % O2 | 193 | 72 |
| g | SRS+13a | pMSC 8 / 1 % O2 | 193 | 75 |
| h | LRS+14 | myoblast secretome | 52 | 29 |
| i | YKK+14 | U373MG glioma cells soluble | 45 | 22 |
| j | YKK+14 | U373MG glioma cells exosome | 56 | 40 |
| k | CRS+15 | MDA-MB-231 breast cancer parental cells | 28 | 25 |
| l | CRS+15 | MDA-BT breast cancer bone tropic cells | 20 | 79 |
| m | RTA+15 | LNCaP and PC3 prostate cancer cell exosomes | 16 | 111 |
| n | RSE+16 | adipose-derived stem cells | 66 | 50 |
| o | CGH+17 | mouse cardiac fibroblasts exosomes | 73 | 71 |
| p | CGH+17 | mouse cardiac fibroblasts secretome | 47 | 75 |
| q | CLY+18 | HCT116 colon cancer cells secretome | 104 | 88 |
| r | DWW+18 | ovarian cancer cell exosomes | 22 | 102 |
| s | FPR+18 | endothelial progenitor cells | 41 | 42 |
| t | ODS+18 | AC10 ventricular cardiomyocyte extracellular vesicles | 14 | 60 |
| u | CWG+19 | U87-MG glioma cell extracellular vesicles | 980 | 1034 |
| v | KAN+19 | cancer-associated fibroblasts secretome | 33 | 26 |
| w | NJVS19 | cancer-associated myofibroblasts | 96 | 54 |
| x | NJVS19 | normal tissue myofibroblasts | 143 | 233 |
| y | PDT+19 | mouse melanoma B16-F0 exosomes | 219 | 988 |

**a**. Tables 2 and 3 of (25). **b**. **c**. Gene names from Supplementary Table S1 of (26), filtered with p-value < 0.05, expression ratio > 1.3 or < 1/1.3 and EF < 2. **d**. Extracted from Supplementary Table SIII of (27): median values of peptide quantification (omitting proteins identified with less than 5 peptides that have different signs of $\log_2$ values); differentially expressed proteins identifed using a $\log_2$ cutoff of 0.2. **e**. Extracted from Table 1 of (28), to include proteins exclusively identified in 1% or 8% $O_2$. **f**. **g**. Extracted from Table 1 of (29), including unique proteins for 1%, 3%, and 8% $O_2$. **h**. GI numbers from Supplementary Data 6 of (30). **i**. **j**. Extracted from Table S1 of (31), to include proteins identified by at least 2 unique peptides and surpassing a $\log_2$ cutoff of 0.5 in soluble or exosome fractions. **k**. **l**. Gene names from Supplementary Information Table 1 of (32), filtered to include proteins with $\log_2$ fold change between air and hypoxia > 0.2 or < -0.2. **m**. Gene names from Supporting Information Tables S1 (normoxic) and S2 (hypoxic) of (33), filtered to include proteins that were exclusively identified in either condition. **n**. Gene names from Tables 1 and 2 of (34). **o**. **p**. Extracted from Tables S2A (exosomes) and S2B (secretome) of (12), keeping proteins with FDR < 0.05. **q**. Supplementary Tables S8-S9 (secretome) of (14). **r**. Supplementary Table 1 of (35). **s**. Table S2 of (36). **t**. Supplementary Material Tables S1 and S2 of (37), filtered to include proteins exclusively identified in hypoxia or normoxia. **u**. Supplementary Tables 1 and 2 of (38), filtered to include proteins uniquely identified in either hypoxia or normoxia. **v**. Proteins identified as up- or down-regulated > 1 SD in Data File S1 of (20) (pooled data from sheets “Soluble Secretome” and “EVs”). **w**. **x**. Extracted from proteinGroups.txt in ProteomeXchange Dataset PXD008104 (39). Expression ratios between hypoxia and normoxia were calculated from LFQ intensity values, and proteins were classified as up- or down-regulated if they had expression ratios > 1.2 or < 1/1.2 in all three experiments for one cell type (CAM or NTM). The Majority protein IDs and mean values of the expression ratios were saved in the data file. **y**. Supplementary Table 2 of (40), filtered with $\log_2$ fold-change cutoff of 1.

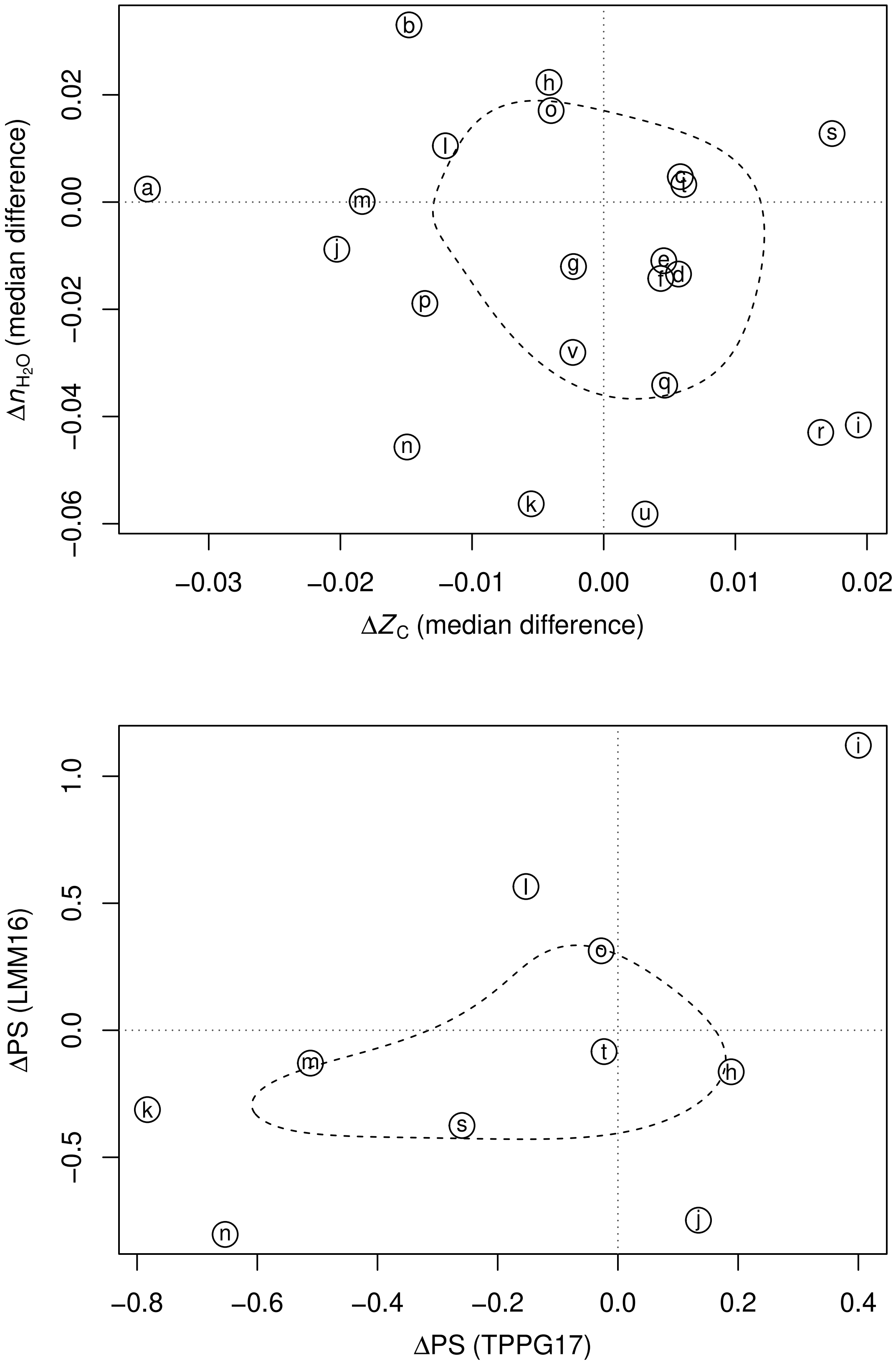


**Fig. S8.** Compositional analysis and phylostrata for hyperosmotic (high salt) experiments in eukaryotic cells.

**Legend for Fig. S8**

Abbreviations: HTS – hypertonic saline; Cmx – cytomix treatment.

| Set | Reference | Description | Down | Up |
|---|---|---|---|---|
| a | DAA+05 | thick ascending limb of Henle's loop cells in 600 (NaCl added) vs 300 mosmol/kg medium | 14 | 24 |
| b | MHN+08 | mouse CGR8 embryonic stem cells in 500 (NaCl added) vs 340 mOsM medium | 23 | 23 |
| c | LTH+11 | *Saccharomyces cerevisiae* Protein in 0.7 M NaCl vs control medium for 30 min | 686 | 203 |
| d | LTH+11 | *Saccharomyces cerevisiae* Protein in 0.7 M NaCl vs control medium for 60 min | 554 | 335 |
| e | LTH+11 | *Saccharomyces cerevisiae* Protein in 0.7 M NaCl vs control medium for 90 min | 537 | 352 |
| f | LTH+11 | *Saccharomyces cerevisiae* Protein in 0.7 M NaCl vs control medium for 120 min | 538 | 351 |
| g | LTH+11 | *Saccharomyces cerevisiae* Protein in 0.7 M NaCl vs control medium for 240 min | 564 | 325 |
| h | OBBH11 | adipose-derived stem cells in 400 mOsm vs 300 mOsm NaCl | 148 | 144 |
| i | LFY+12 | cytoplasm of HEK293 cells in 500 (NaCl added) vs 300 mosmol/kg medium for 1 h | 19 | 36 |
| j | LFY+12 | cytoplasm of HEK293 cells in 500 (NaCl added) vs 300 mosmol/kg medium for 8 h | 20 | 34 |
| k | LFY+12 | cytoplasm of HEK293 cells in 500 (NaCl added) vs 300 mosmol/kg medium for 2 passages | 33 | 66 |
| l | LFY+12 | nucleus of HEK293 cells in 500 (NaCl added) vs 300 mosmol/kg medium for 1 h | 49 | 80 |
| m | LFY+12 | nucleus of HEK293 cells in 500 (NaCl added) vs 300 mosmol/kg medium for 8 h | 39 | 64 |
| n | LFY+12 | nucleus of HEK293 cells in 500 (NaCl added) vs 300 mosmol/kg medium for 2 passages | 22 | 67 |
| o | CLG+15 | human conjunctival epithelial cells in 380 or 480 mOsm vs 280 mOsm NaCl | 25 | 38 |
| p | SCG+15 | *Saccharomyces cervisiae* in 0.4 M NaCl vs control - nodelay | 567 | 266 |
| q | SCG+15 | *Saccharomyces cervisiae* in 0.4 M NaCl vs control - delayed | 219 | 67 |
| r | YDZ+15 | *Yarrowia lipolytica* in 4.21 osmol/kg vs 3.17 osmol/kg NaCl | 14 | 28 |
| s | GAM+16 | human small airway epithelial cells in HTS vs isotonic | 211 | 396 |
| t | GAM+16 | human small airway epithelial cells in HTS.Cmx vs isotonic.Cmx | 303 | 250 |
| u | RBP+16 | *Paracoccidioides lutzii* in 0.1 M KCl vs medium with no added KCl | 160 | 141 |
| v | JBG+18 | *Candida albicans* in 1 M NaCl vs medium with no added NaCl | 84 | 63 |

**a**. Tables II–III of (41). **b**. Table 1 of (42). **c**. **d**. **e**. **f**. **g**. Dataset S3 of (43), filtered to include proteins with $q$-value < 0.05, same direction of change in all 3 replicates for each condition, and median log fold change > 0.2. **h**. Supplementary Table 1 of (44). **i**. **j**. **k**. **l**. **m**. **n**. Supplementary Table S1 of (45) (sheet "All proteins"), filtered to include proteins with $q$-value < 0.1. **o**. Table 2 of (46). **p**. **q**. Supplemental Data S1 of (47) (file "ClusterGroup.AllProteins.051814.csv"), filtered to include proteins in clusters 1–4 (differential expression at all time points (1/4) or after > 20 min delay (2/3)). **r**. Table 1 of (48). **s**. **t**. Supplementary Table 2 of (49), filtered to include proteins with fold change > 2 or < 0.5. **u**. Supplementary Tables 2 and 3 of (50). **v**. Supplementary Tables 1 and 2 of (51).

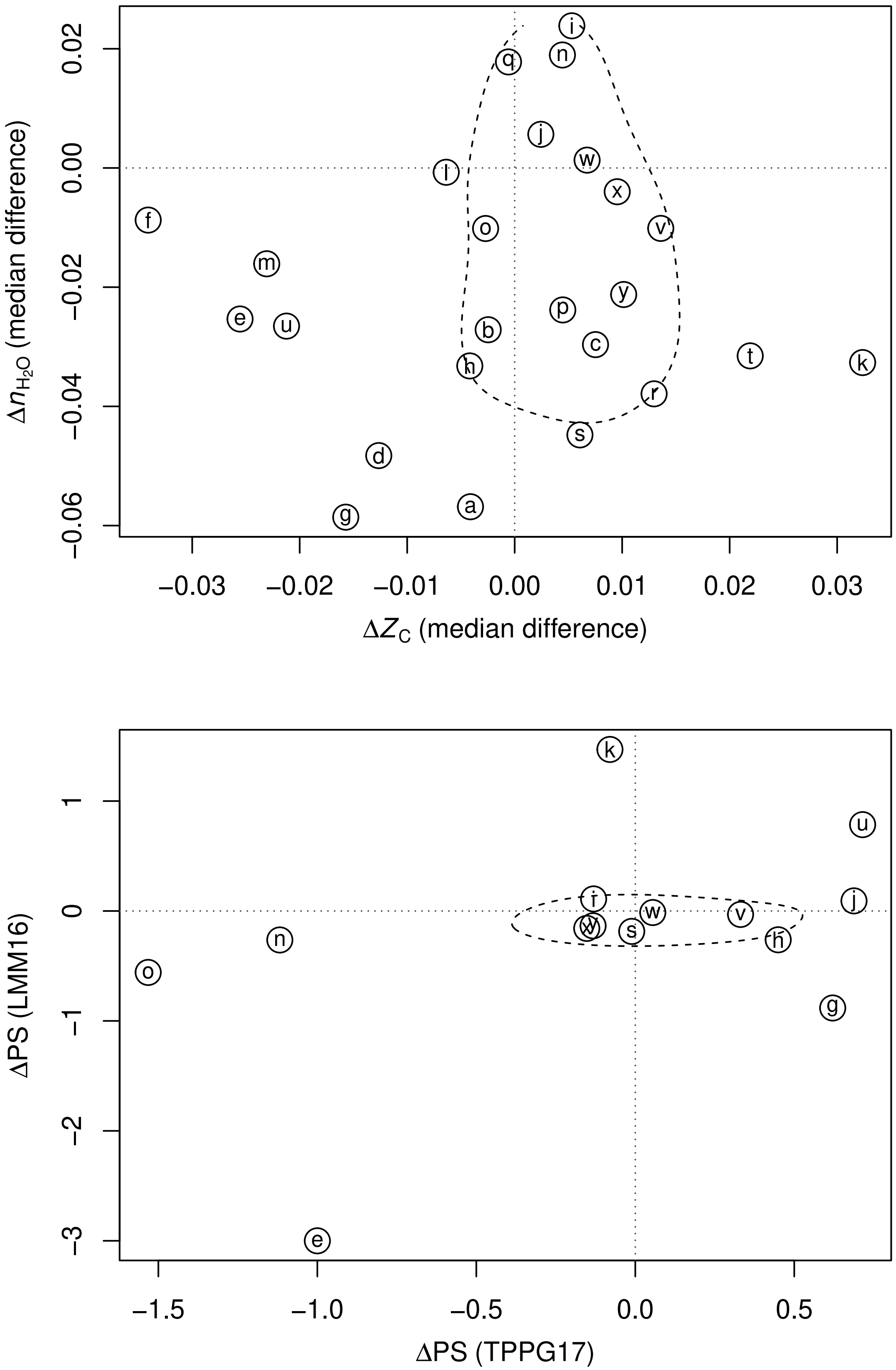


**Fig. S9.** Compositional analysis and phylostrata for hyperosmotic (high glucose) experiments in eukaryotic cells.

#### Legend for Fig. S9

Abbreviations: HUVEC – human umbilical vein endothelial cells; eMP – endothelial-derived microparticles.

| Set | Reference | Description | Down | Up |
|---|---|---|---|---|
| a | PW08 | *Saccharomyces cerevisiae* in very high glucose (300 g/L) vs control (20 g/L) for 2 h | 38 | 44 |
| b | PW08 | *Saccharomyces cerevisiae* in very high glucose (300 g/L) vs control (20 g/L) for 10 h | 33 | 62 |
| c | PW08 | *Saccharomyces cerevisiae* in very high glucose (300 g/L) vs control (20 g/L) for 12 h | 18 | 65 |
| d | WCM+09 | mouse pancreatic islets in 16.7 mM vs 5.6 mM glucose | 63 | 94 |
| e | WFSL09 | bovine aortal endothelial cells in 22 mM vs 5 mM glucose | 67 | 46 |
| f | MFD+10 | rat INS-1E cells in 25 mM vs 11 mM glucose | 51 | 20 |
| g | CCC+12 | retinal pigmented epithelium in 25 mM glucose vs 5.5 mM glucose | 17 | 11 |
| h | CCC+12 | retinal pigmented epithelium in 100 mM glucose vs 5.5 mM glucose | 21 | 24 |
| i | SFG+12 | human pancreatic islets in 15 mM vs 5 mM glucose | 34 | 57 |
| j | CCCC13 | Chang liver cells in 25 mM vs 5.5 mM glucose | 32 | 39 |
| k | CCCC13 | Chang liver cells in 100 mM vs 5.5 mM glucose | 19 | 50 |
| l | CCW+13 | rat INS-1beta cells in 27 mM vs 11 mM glucose | 126 | 60 |
| m | LDB+15 | Chinese hamster ovary cells in 15 g/L vs 5 g/L glucose | 294 | 205 |
| n | BTX+17 | HUVEC eMPs in 5.6 mmol/l glucose + 19.4 mmol/l D-glucose vs 5.6 mmol/l glucose | 28 | 333 |
| o | BTX+17 | HUVEC eMPs in 5.6 mmol/l glucose + 19.4 mmol/l L-glucose vs 5.6 mmol/l glucose | 24 | 377 |
| p | SFKD17 | secretome of murine islets of Langerhans in 25 mM vs 11 mM glucose for 1 day | 59 | 54 |
| q | SFKD17 | secretome of murine islets of Langerhans in 25 mM vs 11 mM glucose for 2 days | 82 | 104 |
| r | HGC+18 | *Lactobacillus casei* BL23 in hyper-concentrated vs isotonic sweet whey | 116 | 64 |
| s | IXA+19 | human aortal endothelial cells in 20 mM vs 5 mM glucose | 49 | 331 |
| t | MHP+20 | rat H9c2 cells in 30 mM vs 5 mM glucose | 17 | 20 |
| u | MHP+20 | human embryonic kidney cells in 30 mM vs 5 mM glucose | 127 | 39 |
| v | MPR+20 | human aortic endothelial cells in 3h.high.glucose | 109 | 133 |
| w | MPR+20 | human aortic endothelial cells in 24h.high.glucose | 224 | 98 |
| x | MPR+20 | human aortic endothelial cells in 3h.high.mannitol | 154 | 127 |
| y | MPR+20 | human aortic endothelial cells in 24h.high.mannitol | 178 | 128 |

**a**. **b**. **c**. Supporting Information Table of (52), filtered to include proteins with expression ratios < 0.9 or > 1.1 and with *p*-values < 0.05. **d**. Supplemetary Table ST4 of (53), filtered to include the proteins with ANOVA *p*-value < 0.01 (red- and blue-highlighted rows in the source table), and applying the authors' criterion that proteins be identified by 2 or more unique peptides in at least 4 of the 8 most intense LC-MS/MS runs. **e**. Supplementary Table of (54), filtered to include proteins with fold change > 1.2 or < 0.8. **f**. Table 1 of (55). **g**. **h**. Table 1 of (56). **i**. Proteins identified as differentially abundant in Supporting Information Table S5 of (57), filtered to include proteins with fold change > 2 or < 0.5. **j**. **k**. Table 1 of (58). **l**. Supplementary Table 1 of (59), filtered to include proteins with average fold change > 2.5 or < 0.4. **m**. Supporting Information Table S4 of (60) for up- (Cluster 1) and down- (Cluster 5) regulated proteins. **n**. **o**. Electronic supplementary material Table 1 of (61). **p**. **q**. Supplementary Table S1 of (62), filtered to include proteins with fold change > 2 or < 0.5 (ratios were computed from medians of iBAQ values for three replicates after quantile normalization). **r**. Supplementary Figure 1 of (63). **s**. Supplementary Tables 1 and 2 of (64). **t**. **u**. Supplementary Table 2 of (65) (sheets "H9c2" and "HEK"). **v**. **w**. **x**. **y**. Treatment with high glucose (12.5 mmol/L) or high mannitol (7.0 mmol/L + 5.5 mmol/L glucose) followed by insulin, compared to normal glucose (5.5 mmol/L) followed by insulin. Source: Supplementary Tables S2–S6 of (66) (proteins uniquely identified in treatment and control conditions).

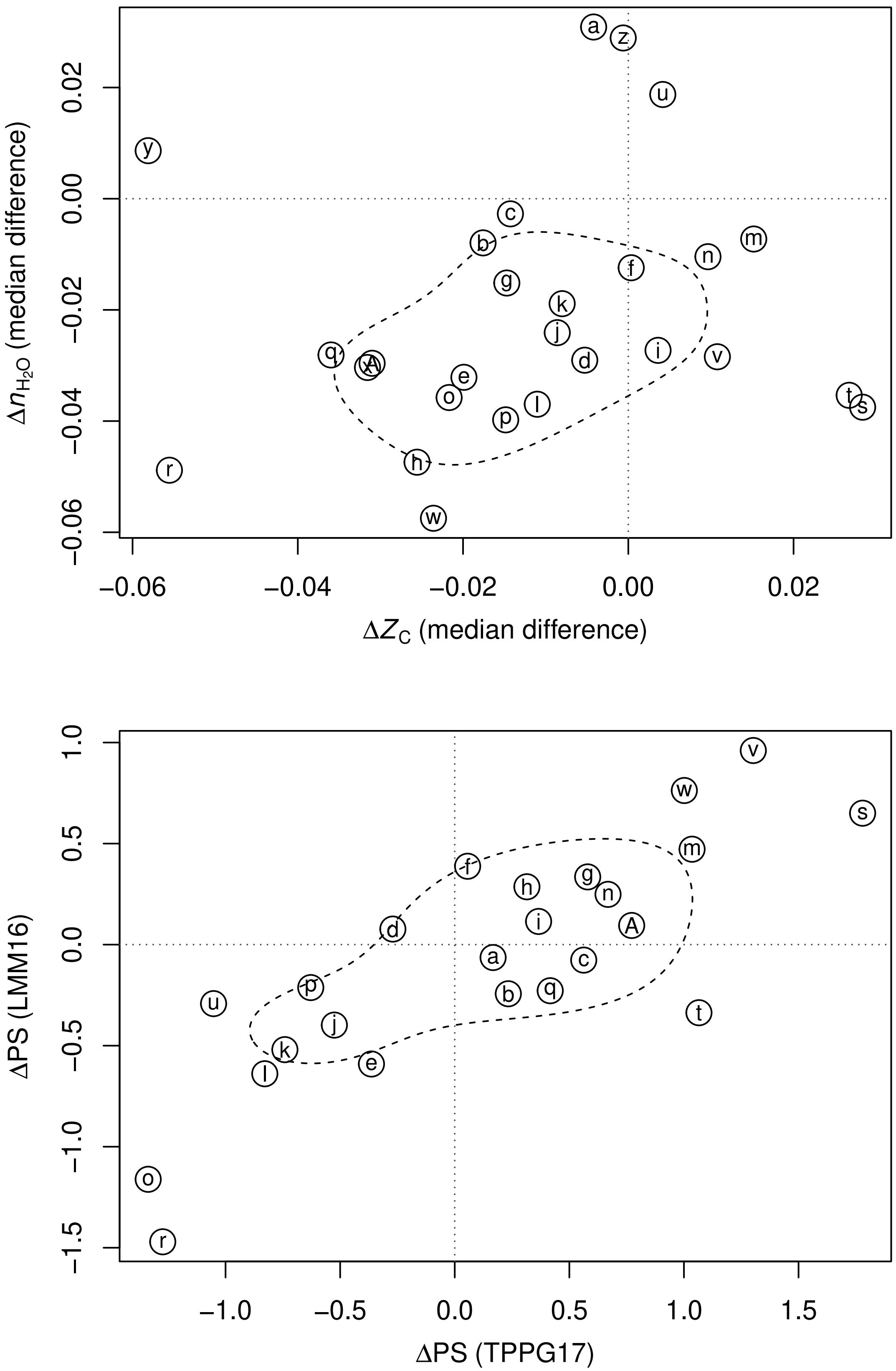


**Fig. S10.** Compositional analysis and phylostrata for 3D cell culture vs 2D cell culture experiments.

### Legend for Fig. S10

Abbreviations: P2 – passage 2; P5 – passage 5; MSC – mesenchymal stromal cells.

| Set | Reference | Description | Down | Up |
|---|---|---|---|---|
| a | PLC+10 | HepG2 hepatocellular carcinoma cells | 35 | 47 |
| b | MHG+12 | MCF-7 breast cancer cells P5 | 409 | 337 |
| c | MHG+12 | MCF-7 breast cancer cells P2 | 248 | 214 |
| d | MVC+12 | HT29 colon cancer cells perinecrotic | 48 | 52 |
| e | MVC+12 | HT29 colon cancer cells necrotic | 101 | 186 |
| f | YYW+13 | HNSCC tumor spheres | 58 | 65 |
| g | ZMH+13 | HUVEC Matrigel 12h | 48 | 58 |
| h | ZMH+13 | HUVEC Matrigel 24h | 196 | 239 |
| i | HKX+14 | U251 glioma cells | 295 | 68 |
| j | KDS+14 | hESC spheroids | 270 | 52 |
| k | KDS+14 | hiPSC spheroids | 422 | 53 |
| l | KDS+14 | hPSC spheroids | 612 | 98 |
| m | RKP+14 | colorectal cancer-derived cells | 93 | 174 |
| n | SAS+14 | SK-N-BE2 neuroblastoma spheroids | 70 | 116 |
| o | WRK+14 | HepG2/C3A hepatocellular carcinoma | 125 | 291 |
| p | MTK+15 | OV-90AD ovarian cancer multicellular aggregates | 39 | 90 |
| q | YLW+16 | HT29 colon carcinoma | 116 | 225 |
| r | KJK+18 | SW480 colorectal cancer | 247 | 136 |
| s | TGD18 | normal human skin fibroblasts | 15 | 57 |
| t | TGD18 | cancer-associated fibroblasts | 43 | 90 |
| u | EWK+19 | glioblastoma spheroids | 110 | 171 |
| v | GADS19 | skin fibroblasts | 94 | 49 |
| w | HLC19 | HepG2 hepatocellular carcinoma cells | 573 | 590 |
| x | LPK+19 | mouse 3T3-L1 preadipocytes | 76 | 105 |
| y | LPK+19 | mouse 3T3-L1 adipocytes | 97 | 208 |
| z | LPK+19 | mouse 3T3-L1 macrophages | 220 | 134 |
| A | DKM+20 | bone marrow-derived MSCs aggregates | 57 | 117 |

**a**. Gene names from Supporting Information Table 1S of (67). **b**. **c**. Sheets 2 and 3 in Table S1 of (68). **d**. **e**. Supplemental Table 1C of (69), filtered to include proteins with expression ratios < 0.77 or > 1.3. **f**. Supplemental Table S5 of (70). **g**. **h**. Supplemental Table S4 of (71). **i**. Table S1 of (72). **j**. **k**. **l**. Supplemental Table S1 of (73) (hESC: human embryonic stem cells; hiPSC: human induced pluripotent stem cells; hPSC: human pluripotent stem cells). **m**. Table S2 of (74), filtered to include proteins that have differences in spectral counts recorded in at least two of three experiments, absolute overall fold change is ≥ 1.5 or ≤ 2/3, and *p*-value is < 0.05. **n**. Supplementary Figure 2 of (75). **o**. P1_Data sheet in the Supporting Information file of (76). **p**. Supplemental Table S1 of (77), filtered to exclude marked contaminants and reverse sequences and to include proteins with "Ratio H/L normalized" > 1.5 or < 2/3. **q**. Tables S1a and S1b of (78). **r**. Tables S2 and S3 of (79). **s**. **t**. Supplementary Table 3 of (80). **u**. Extracted from proteinGroups.txt in ProteomeXchange Dataset PXD008244/txt.zip [(81)], including proteins quantified in at least two replicates in each condition (adherent and spheroid) and with median fold change > 2 or < 0.5. **v**. Extracted from Supplementary Table 1 of (82). Values were quantile normalized, then ratios were calculated between 3D and 2D cultures for each treatment (control, CBD, UVA, UVA+CBD, UVB, UVB+CBD). Ratios > 1.2 or < 1/1.2 in at least 4 treatments were used to identify differentially expressed proteins. **w**. Supplementary Data I, sheet "Volcano plot 2Dv3D" of (83). **x**. **y**. **z**. Supporting Table 1 of (84), sheets "3P2P" (mono-cultured preadipocytes), "3A2A" (mono-cultured adipocytes), "3C2C" (co-cultured adipocytes with macrophages), filtered to include proteins marked as "T-test Significant" and with absolute value of "N: T-test Difference" > $\log_2(0.5)$. **A**. Supplementary Table S2 of (85), filtered to include proteins with fold change > 2 or < 0.5 for all donors.

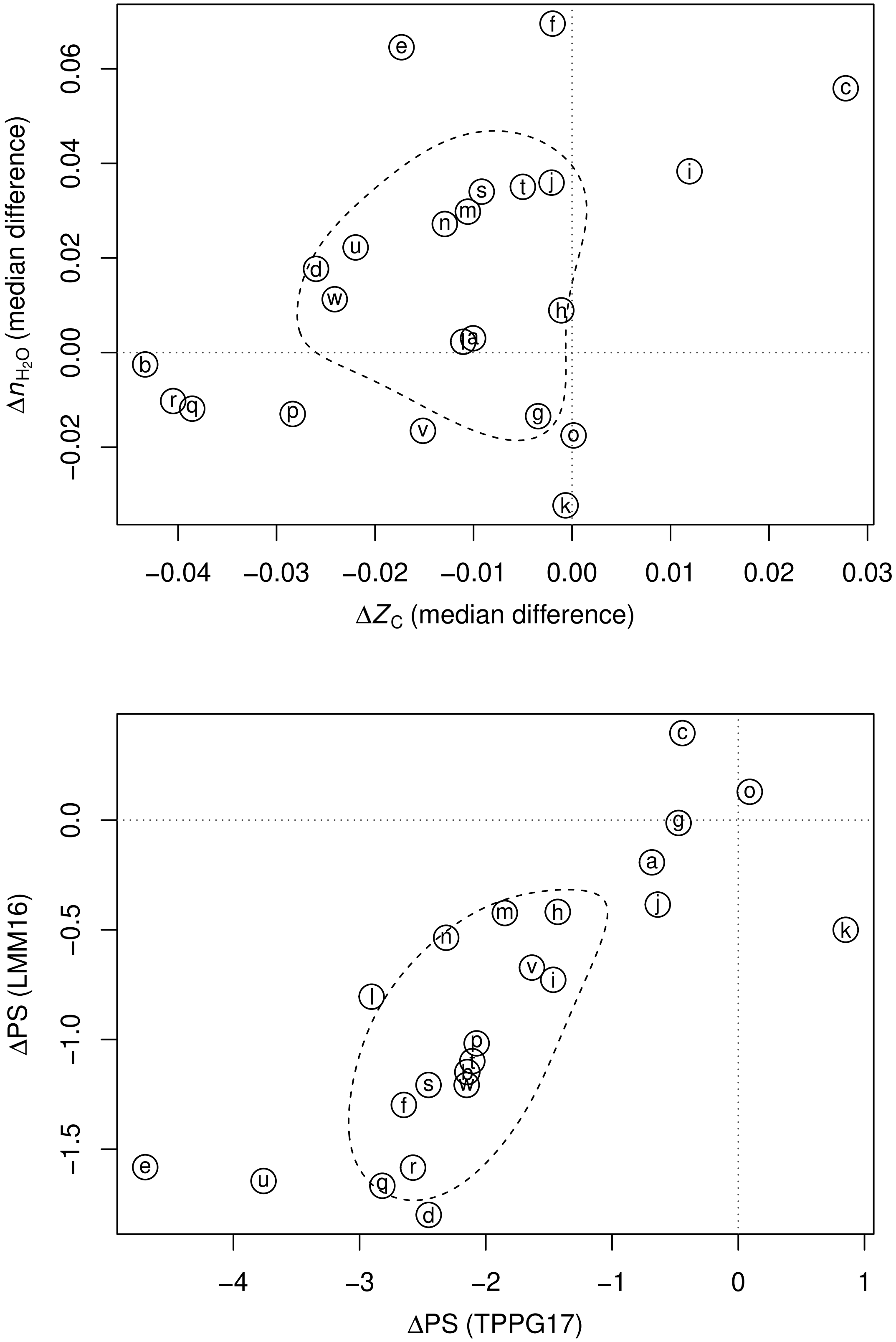


**Fig. S11.** Compositional analysis and phylostrata for differentially expressed proteins in breast cancer.

### Legend for Fig. S11

Abbreviations: T – tumor; N – normal; LCM – laser capture microdisecction; FFPE – formalin-fixed paraffin-embedded; IDC – infiltrating ductal carcinoma; DCIS – ductal carcinoma in situ; TNBC – triple-negative breast cancer.

| Set | Reference | Description | Down | Up |
|---|---|---|---|---|
| a | AMG+08 | T / periphery | 56 | 128 |
| b | CIR+10 | LCM T / N | 59 | 62 |
| c | SRG+10 | invasive carcinoma / N | 56 | 61 |
| d | HTP+11 | microvessels IDC / nonmalignant | 40 | 86 |
| e | GTM+12 | IDC / benign | 27 | 166 |
| f | GTM+12 | IDC / adjacent N | 71 | 122 |
| g | LLL+13 | TNBC tumor / paraneoplastic | 128 | 80 |
| h | SRS+13 | DCIS / matched N | 56 | 50 |
| i | SRS+13 | IDC / matched N | 73 | 71 |
| j | GSB+14 | T / distant N | 88 | 108 |
| k | PPH+14 | multiple subtypes T / N | 54 | 40 |
| l | CVJ+15 | TNBC T / N | 92 | 124 |
| m | PGT+16 | early stage T / N | 21 | 38 |
| n | PGT+16 | late stage T / N | 24 | 42 |
| o | PBR+16 | FFPE T / adjacent N | 406 | 563 |
| p | BST+17 | tumor epithelium / N | 234 | 245 |
| q | TZD+18 | T / adjacent N all | 268 | 1836 |
| r | TZD+18 | T / adjacent N basal | 291 | 1058 |
| s | GCS+19 | T / contralateral N | 162 | 235 |
| t | GCS+19 | T / adjacent N | 159 | 246 |
| u | LLC+19 | T / adjacent N | 93 | 48 |
| v | LLF+20 | TNBC grade I.II T / adjacent N | 556 | 309 |
| w | LLF+20 | TNBC grade III T / adjacent N | 667 | 359 |

**a**. Extracted from Supporting Information RTF files of (86). Proteins identified by any number of peptides in both cancer and matched periphery were excluded; of the remaining proteins those identified by at least two peptides were used. **b**. Table S2(a) of (87) (proteins differentially abundant at or above 99% confidence level). **c**. Table 4 of (88). **d**. Tables 1 and 2 of (89). **e**. **f**. Supporting Table 6 of (90). **g**. Table S1 of (91). **h**. **i**. Table 2 of (92). For DCIS (3 patients), proteins were classified as up/down regulated if 2 or more ratios were greater/less than 1, and no ratios were less/greater than 1. For IC (4 patients), proteins were classified as up/down regulated if 3 or more ratios were greater/less than 1, and no ratios were less/greater than 1. **j**. Extracted from Table S2 of (93). Values in all LFQ columns (distant, near, tumor) were quantile normalized, then the ratio between tumor and distant was calculated. Proteins with normalized LFQ ratios > 1.2 or < 1/1.2 and $p$-value < 0.05 were identified as differentially expressed. **k**. Supplementary Tables 12, 13, 14 and 15 of (94), filtered to include proteins that are up- or down-regulated in all subtypes (LUM, LUMHER, HER, TN). **l**. Supplemental Data S3 of (95), filtered to include proteins with min and max credible intervals for expression ratios that are both <1 or >1. **m**. **n**. Supplementary Table S3 of (96). **o**. Table S4A of (97). **p**. Supplemental Table 4 of (98), filtered to include proteins with $p$-value < 2. **q**. **r**. Tables S5-1 (differentially expressed proteins for 52 tumor / non-cancerous tissue pairs) and S5-2 (13 basal-like tumor / non-cancerous tissue pairs) of (99), filtered to include proteins with $\log_2$ fold change > 1 or < -1. **s**. **t**. Supplementary File S1 of (100) (PT: primary breast tissue; NCT: non tumor contralateral breast tissue; ANT: non tumor adjacent breast tissue). **u**. Supplementary Table 2 of (101). **v**. **w**. Supplementary Tables S2 and S3 of (102).

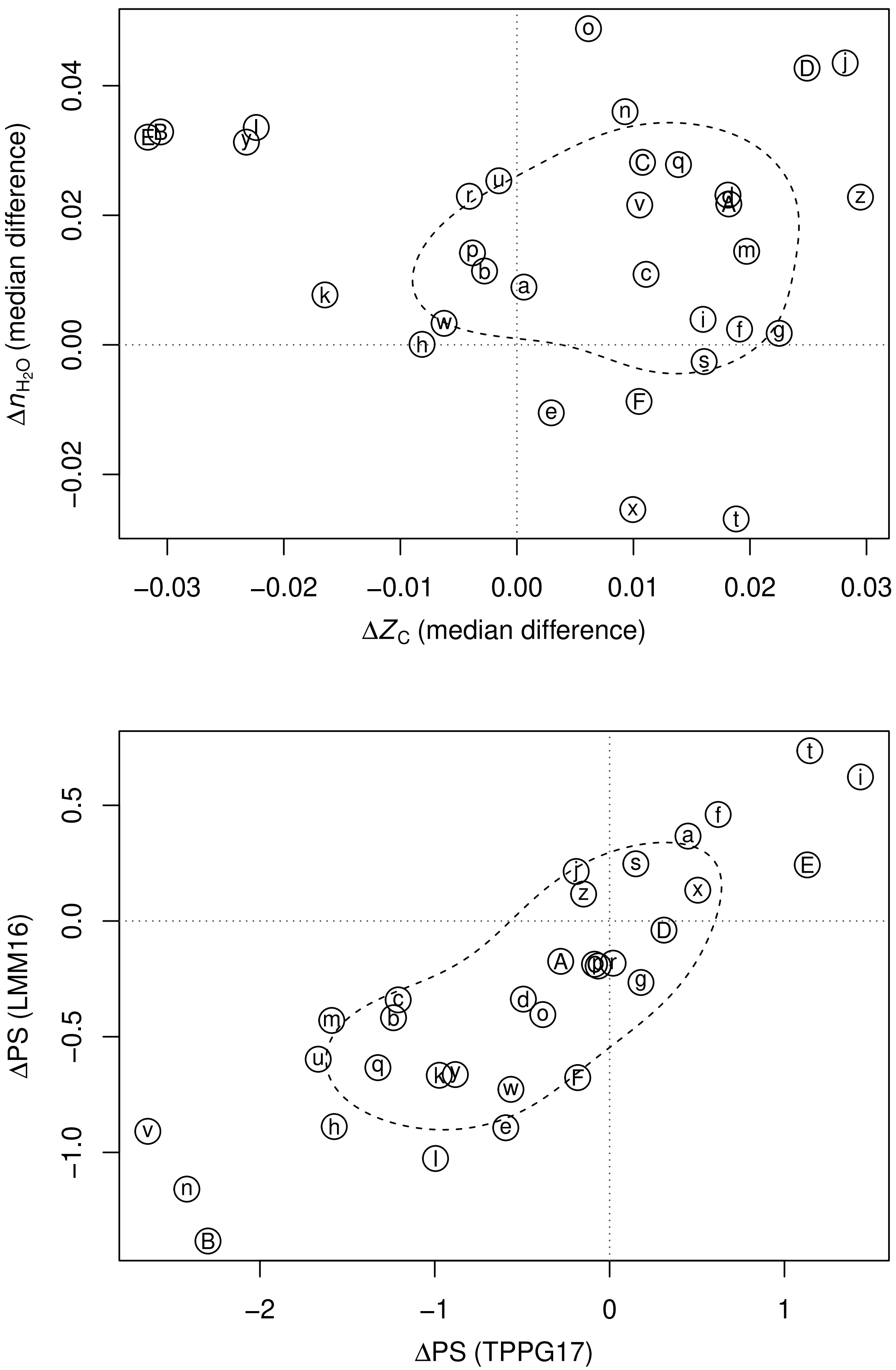


**Fig. S12.** Compositional analysis and phylostrata for differentially expressed proteins in colorectal cancer.

### Legend for Fig. S12

Abbreviations: T – tumor; N – normal; LCM – laser capture microdisecction; FFPE – formalin-fixed paraffin-embedded; MSS – microsatellite stable; CIS – carcinoma in situ; ICC – invasive colonic carcinoma.

| Set | Reference | Description | Down | Up |
|---|---|---|---|---|
| a | WTK+08 | T / N | 57 | 70 |
| b | XZC+10 | stage I / normal | 48 | 166 |
| c | XZC+10 | stage II / normal | 77 | 321 |
| d | ZYS+10 | microdissected T / N | 60 | 57 |
| e | BPV+11 | stage I / normal | 109 | 72 |
| f | BPV+11 | stage II / normal | 164 | 140 |
| g | BPV+11 | stage III / normal | 63 | 131 |
| h | BPV+11 | stage IV / normal | 42 | 26 |
| i | JCF+11 | T / N | 72 | 45 |
| j | MRK+11 | adenocarcinoma / normal | 350 | 232 |
| k | SHHS11 | LCM T / N | 28 | 43 |
| l | FGW+12 | T / matched N | 48 | 34 |
| m | KYK+12 | MSS-type T / N | 73 | 175 |
| n | WOD+12 | LCM FFPE T / adjacent N | 79 | 677 |
| o | CZD+14 | T / N | 52 | 74 |
| p | STK+15 | membrane enriched T / N | 113 | 66 |
| q | WDO+15 | LCM FFPE T / adjacent N | 879 | 1281 |
| r | LXM+16 | biopsy T / N | 191 | 178 |
| s | PHL+16 | CIS / N | 169 | 138 |
| t | PHL+16 | ICC / N | 129 | 100 |
| u | CTW+17 | organoid T / N | 227 | 78 |
| v | HZW+17 | T / adjacent N | 126 | 589 |
| w | LLL+17 | LCM cancer / non-neoplastic mucosa | 110 | 77 |
| x | NKG+17 | T / N | 48 | 77 |
| y | QMB+17 | FFPE T / N | 25 | 53 |
| z | TMS+17 | epithelial T / N | 158 | 166 |
| A | ZLY+17 | T / N | 58 | 54 |
| B | AKG+18 | T / N | 43 | 426 |
| C | STA+19 | non-metastatic colon cancer, T / N | 437 | 317 |
| D | STA+19 | metastatic colon cancer, T / N | 436 | 356 |
| E | VHW+19 | T / adjacent N | 417 | 31 |
| F | WYL+19 | tumor-associated / normal vascular endothelial cells | 97 | 119 |

**a**. Table 1 and Supplementary Data 1 of (103) (Swiss-Prot and UniProt accession numbers from Supplementary Data 2). **b**. **c**. IPI accession numbers from Supplemental Table 4 of (104). **d**. IPI accession numbers from Supplemental Table 4 of (105). **e**. **f**. **g**. **h**. Gene names from supplemental Table 9 of (106). **i**. Supplementary Table 2 of (107). **j**. Table S8 of (108). **k**. Supplementary Table 1 of (109). **l**. Appendix of (110). **m**. Gene names from Supplementary Table 4 of (111), filtered to include proteins with expression ratio > 2 or < 0.5 in both mTRAQ and cICAT analyses. **n**. Supplementary Table 4 of (112). **o**. Table 2 of (113). **p**. Ensembl protein IDs from Supporting Table 2 of (114). **q**. Proteins marked as having a significant change between normal tissue (N) and adenocarcinoma (C) in SI Table 3 of (115). **r**. SI Table S3 of (116), filtered to include proteins with $p$-value < 0.05. **s**. **t**. Gene names from Supplementary Table 4 of (117), for differential expression between normal colonic mucosa (NC) and carcinoma in situ (CIS) or invasive colorectal cancer (ICC). **u**. Table S3 of (118). **v**. Dataset 6A of (119). **w**. Supplementary Material Table S1 of (120). **x**. Table 2 of (121). **y**. Table S2 of (122), filtered to include comparisons between adenocarcinoma and diverticular disease. **z**. Table S1 of (123), filtered to include proteins that are consistently up- or down-regulated in at least 11 of 12 patients. **A**. IPI accession numbers from Table S2 of (124). **B**. Supplementary Table 1B of (125), filtered to include proteins with expression ratio > 3/2 or < 2/3. **C**. **D**. Supplementary Table S2 of (126), filtered to include proteins with $\log_2$ ratio > ± the standard deviation of values for all quantified proteins. **E**. Online data from (127) (file: Human__CPTAC_COAD__PNNL__Proteome__TMT__03_01_2017__BCM__Gene__Tumor_Normal_log2FC.cct), filtered to include proteins with median $\log_2$ ratio > 1 or < -1. **F**. Supplementary Information Table S2 of (128).

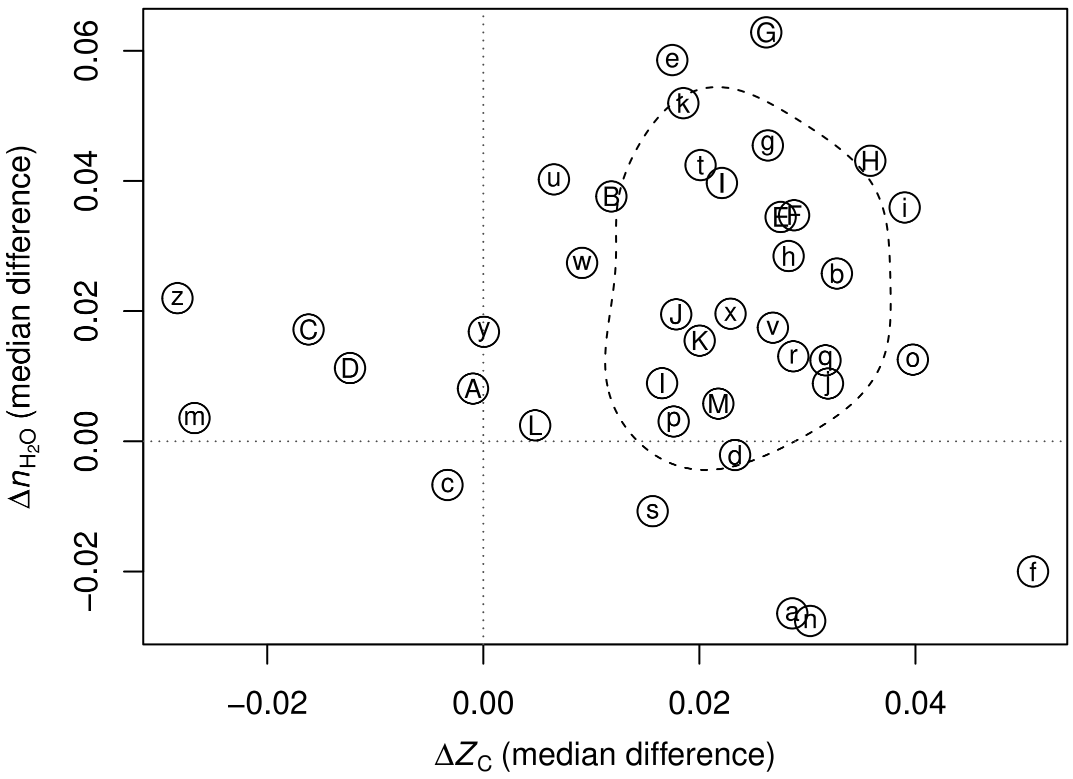


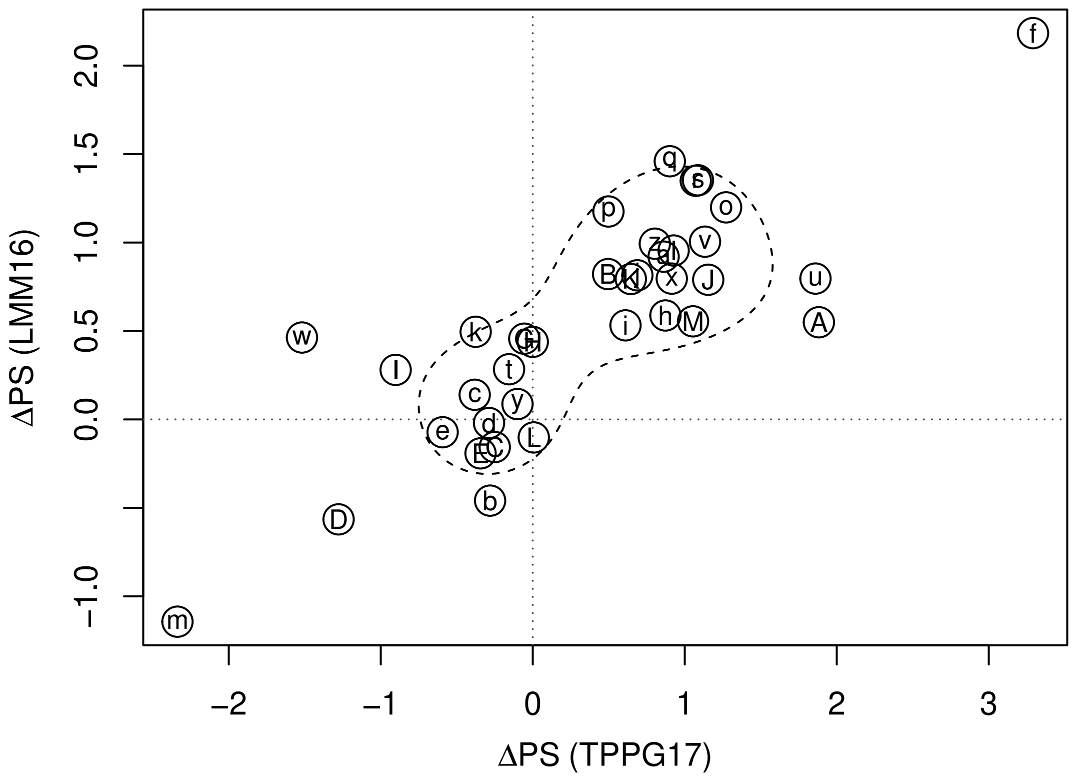


**Fig. S13.** Compositional analysis and phylostrata for differentially expressed proteins in liver cancer.

### Legend for Fig. S13

Abbreviations: EGF – epidermal growth factor; G1, G2, G3 – well-differentiated, moderately differentiated, poorly differentiated tumors; T1, T2, T3 – T1N0M0, T2N0M0, T3N0M0 tumor-node-metastasis classification.

| Set | Reference | Description | Down | Up |
|---|---|---|---|---|
| a | LHT+04 | T / N | 79 | 44 |
| b | BLP+05 | T / N | 61 | 22 |
| c | LTZ+05 | T / N | 35 | 39 |
| d | DTS+07 | T / N | 20 | 23 |
| e | SXS+07 | T / N | 46 | 21 |
| f | CHN+08 | T / N | 93 | 58 |
| g | RLA+10 | T / N rat transitional endoplasmic reticulum | 130 | 135 |
| h | LMG+11 | T / N nuclear | 411 | 264 |
| i | LMG+11 | T / N cytoskeletal | 280 | 273 |
| j | LRL+12 | T / N | 102 | 161 |
| k | KOK+13 | T / N | 53 | 38 |
| l | MBK+13 | T / N | 192 | 280 |
| m | XWS+14 | T / N | 135 | 504 |
| n | BSG15 | T / N EGF transgenic mice | 34 | 71 |
| o | RPM+15 | T / N | 131 | 479 |
| p | NBM+16 | T / N G1 | 42 | 80 |
| q | NBM+16 | T / N G2 | 81 | 82 |
| r | NBM+16 | T / N G3 | 237 | 556 |
| s | NMB+16 | T / N | 121 | 426 |
| t | QXC+16 | T / N T1 | 198 | 132 |
| u | QXC+16 | T / N T2 | 193 | 172 |
| v | QXC+16 | T / N T3 | 202 | 185 |
| w | GJZ+17 | T / N | 26 | 25 |
| x | GWS+17 | T / N | 145 | 191 |
| y | QPP+17 | T / N | 121 | 72 |
| z | WLL+17 | T / N small | 48 | 31 |
| A | WLL+17 | T / N medium | 21 | 39 |
| B | WLL+17 | T / N large | 39 | 63 |
| C | WLL+17 | T / N huge | 60 | 129 |
| D | BOK+18 | T / N | 108 | 59 |
| E | YXZ+18 | T / N | 145 | 111 |
| F | BEM+20 | T / N mouse | 93 | 138 |
| G | GZD+19 | T / N protein | 403 | 52 |
| H | GZD+19 | T / N phosphoprotein | 213 | 68 |
| I | JSZ+19 | T / N | 359 | 1649 |
| J | ZZL+19 | T / N | 162 | 173 |
| K | GZL+20 | T / N | 234 | 769 |
| L | SCL+20 | T / N mitochondrial differential | 38 | 43 |
| M | SCL+20 | T / N mitochondrial unique | 419 | 966 |

**a**. Table III of (129). **b**. Tables 2 and 3 of (130). **c**. Table 2 of (131). **d**. Table 1 of (132), including proteins identified in either tumor homogenates or laser microdissected samples. **e**. Supplemental Table S1 of (133). **f**. Tables 1–3 of (134). **g**. Supplemental Tables S3A and S3C of (135), filtered to include proteins with *p*-value $< 0.05$. **h**. **i**. IPI numbers from Supplemental Tables S3 (nuclear proteins) and S4 (cytoskeletal proteins) of (136), filtered to include proteins with median fold-change > 2 or < 0.5. **j**. Supplemental Table 5 of (137). **k**. Supplementary Table 3 of (138). **l**. Supplemental Data S5 (sheet "LF_proteins") of (139). **m**. Supporting Information SI-S2 of (140). **n**. Table 1 of (141). **o**. Supplementary Table S1 of (142), filtered to include proteins with > 1 peptide used for quantification and fold change > 2 in either direction. **p**. **q**. **r**. Supplementary Data Table S2 of (143) (sheets "G1 vs C", "G2 vs C", and "G3 vs C"), filtered to include proteins with *p*-value < 0.05, quantified in at least half of both tumor and control samples, and median $\log_2$ fold change > 1 or < -1. **s**. Supporting Information Table S-4 (sheet "Filtered protein list") of (144). **t**. **u**. **v**. Supplementary Tables S3–S5 of (145). **w**. Table 4 of (146).

**x**. Supplementary Table S5 of (147). **y**. Supplementary Table 5 of (148), filtered to include proteins with median fold change > 2 or < 0.5 (UniProt IDs from Supplementary Table 1). **z**. **A**. **B**. **C**. Supplementary Table S3 of (149), filtered to include proteins with *p*-value < 0.05 and fold change > 2 or < 0.5. **D**. Supplemental Table S2 of (150) (data for tumor vs. peritumor), filtered to include proteins with *q*-value < 0.05, quantified in at least 3 tumors, same direction of change in all tumors, and median $\log_2$ fold change > 1 or < -1. **E**. Supplementary Table S2 of (151). **F**. Dataset of (152), filtered to include proteins quantified in more than half of each of control and cancer samples and median fold change > 2 or < 0.5. **G**. **H**. Supplemental Table S3 of (153) (sheets "1. 1,274 DF proteins" and "2. 859 DF phosphoproteins"), filtered to include proteins with $\log_2$ fold change > 1 or < -1. **I**. Supplementary Table 6 of (154). **J**. Supporting Information Table S5 of (155) (sheet "c_proteins"), filtered to include proteins quantified in at least half of tumor and normal samples and with fold change > 2 or < 0.5. **K**. Supporting Information Table S4 of (156) (sheet "B_Molecule"). **L**. **M**. Supplementary Tables S1–S2 (proteins with differential expression between non-tumor and tumor regions) and S3–S4 (all proteins identified in each region) of (157). Common accession numbers in Tables S3 and S4 were eliminated to yield proteins uniquely identified in either tumor or non-tumor regions.

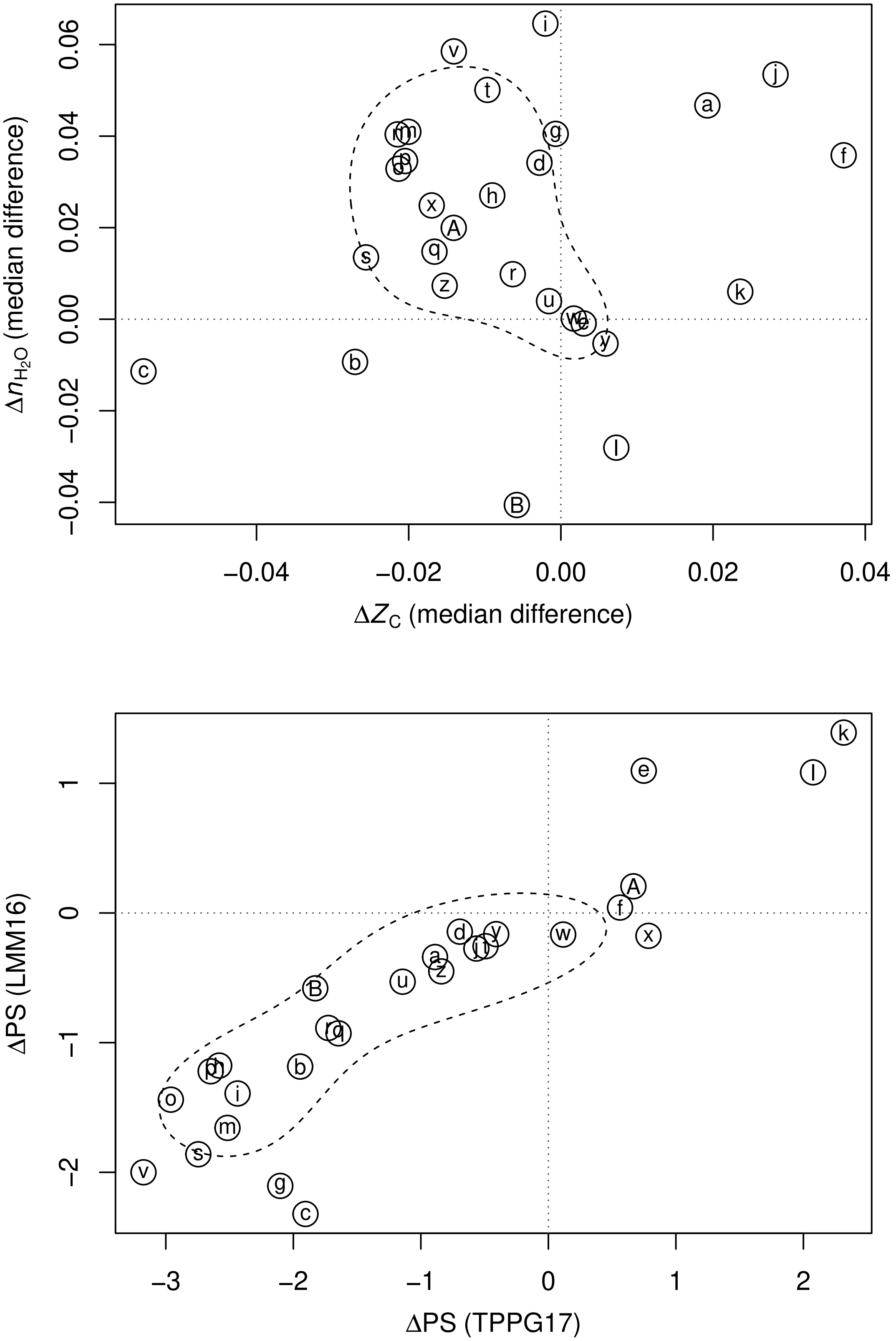


**Fig. S14.** Compositional analysis and phylostrata for differentially expressed proteins in lung cancer.

### Legend for Fig. S14

Abbreviations: NSCLC – non-small cell lung cancer; SCC – squamous cell carcinoma; ADC – adenocarcinoma; ANT – adjacent or patient-matched normal (non-malignant, benign) tissue; NBE – normal bronchial epithelium; LCM – laser capture microdissection; FFPE – formalin-fixed paraffin-embedded.

| Set | Reference | Description | Down | Up |
|---|---|---|---|---|
| a | LXC+06 | SCC / NBE | 22 | 44 |
| b | KHA+12 | ADC / pooled normal | 25 | 25 |
| c | KHA+12 | SCC / pooled normal | 23 | 24 |
| d | YLL+12 | LCM SCC / NBE | 47 | 46 |
| e | ZZD+12 | LCM LSCC / NBE | 54 | 41 |
| f | ZZY+13 | plasma membrane ADC / ANT | 24 | 21 |
| g | LLY+14 | SCC / normal | 29 | 48 |
| h | LWT+14 | NSCLC / ANT | 345 | 1240 |
| i | ZLH+14 | membrane microdissected ADC / ANT | 310 | 257 |
| j | ZLS+14 | endothelial SCC / normal | 61 | 24 |
| k | KNT+15 | FFPE LPIA / pseudo-normal | 346 | 66 |
| l | BLL+16 | NSCLC proteome | 354 | 240 |
| m | FGP+16 | adenocarcinoma / ANT | 81 | 285 |
| n | JCP+16 | mouse endothelial tumor / normal | 18 | 28 |
| o | HHH+16 | pN0 / normal | 210 | 135 |
| p | HHH+16 | pN1 / normal | 233 | 170 |
| q | HHH+16 | pN2.M1 / normal | 154 | 158 |
| r | TLB+16 | NSCLC / ANT | 346 | 1059 |
| s | FGW+17 | adenocarcinoma / ANT | 71 | 1031 |
| t | LZW+17 | mitochondria-related proteins adenocarcinoma / normal | 30 | 30 |
| u | SFS+17 | SCC / ANT LF | 422 | 311 |
| v | WLC+17 | NSCLC / ANT | 20 | 71 |
| w | YCC+17 | SCC.Oncogene / ANT | 280 | 115 |
| x | YCC+17 | SCC.TSG / ANT | 207 | 69 |
| y | YCC+17 | SCC.Glycoprotein / ANT | 64 | 86 |
| z | YCC+17 | ADC.Oncogene / ANT | 385 | 72 |
| A | YCC+17 | ADC.TSG / ANT | 286 | 42 |
| B | YCC+17 | ADC.Glycoprotein / ANT | 83 | 134 |

**a**. Table 1 of (158). **b**. **c**. Gene names from Tables II and III of (159). **d**. Gene names from Table 2 of (160). **e**. IPI numbers from Table 1 of (161). **f**. Table 2 of (162). **g**. Table 1 of (163). **h**. Supplementary data 1b of (164). **i**. Supplementary Data of (165). **j**. Gene names from Supporting Information Table S5 of (166) (differential expression between normal endothelial cells and both paratumor and tumor endothelial cells). **k**. Table S3 of (167) (LPIA: lepidic predominant invasive adenocarcinoma vs pN: pseudo-normal), filtered to include proteins with $\log_2$ fold change > 1 or < -1 and *p*-value < 0.05. **l**. UniProt names from Table S2 of (168), filtered to include proteins with $\log_2$ fold change > 1.5 or < -1/1.5 and *p*-value < 0.05. **m**. Gene names from Table S2 of (169). **n**. Gene names from Table 1 of (170). **o**. **p**. **q**. Supplemental Table S3B of (171) (primary tumor / normal comparison; pN0: no nodes involved; pN1: ipsilateral peribronchial/interlobar/hilar LN metastasis; pN2: ipsilateral mediastinal LN metastasis; M1: distant metastasis or malignant effusion). **r**. Supplemental Table 2 of (172), filtered with adjusted *p*-value < 0.05 and fold change ≥ 1.5 or ≤ 2/3. **s**. Gene names from Supplementary Table S1 of (173), filtered with *p*-value < 0.05. **t**. Table 1 of (174). **u**. Supplementary Table S2 of (175) (sheet LF: label-free quantification). **v**. Supplementary Table 1 of (176). **w**. **x**. **y**. **z**. **A**. **B**. Gene names from Tables S2–S6 of (177) filtered to keep proteins with fold change ≥1.5 or ≤2/3 (Oncogene: oncogene-coded proteins; TSG: tumor suppressor gene-coded proteins; Glycoprotein: glycoproteomics data).

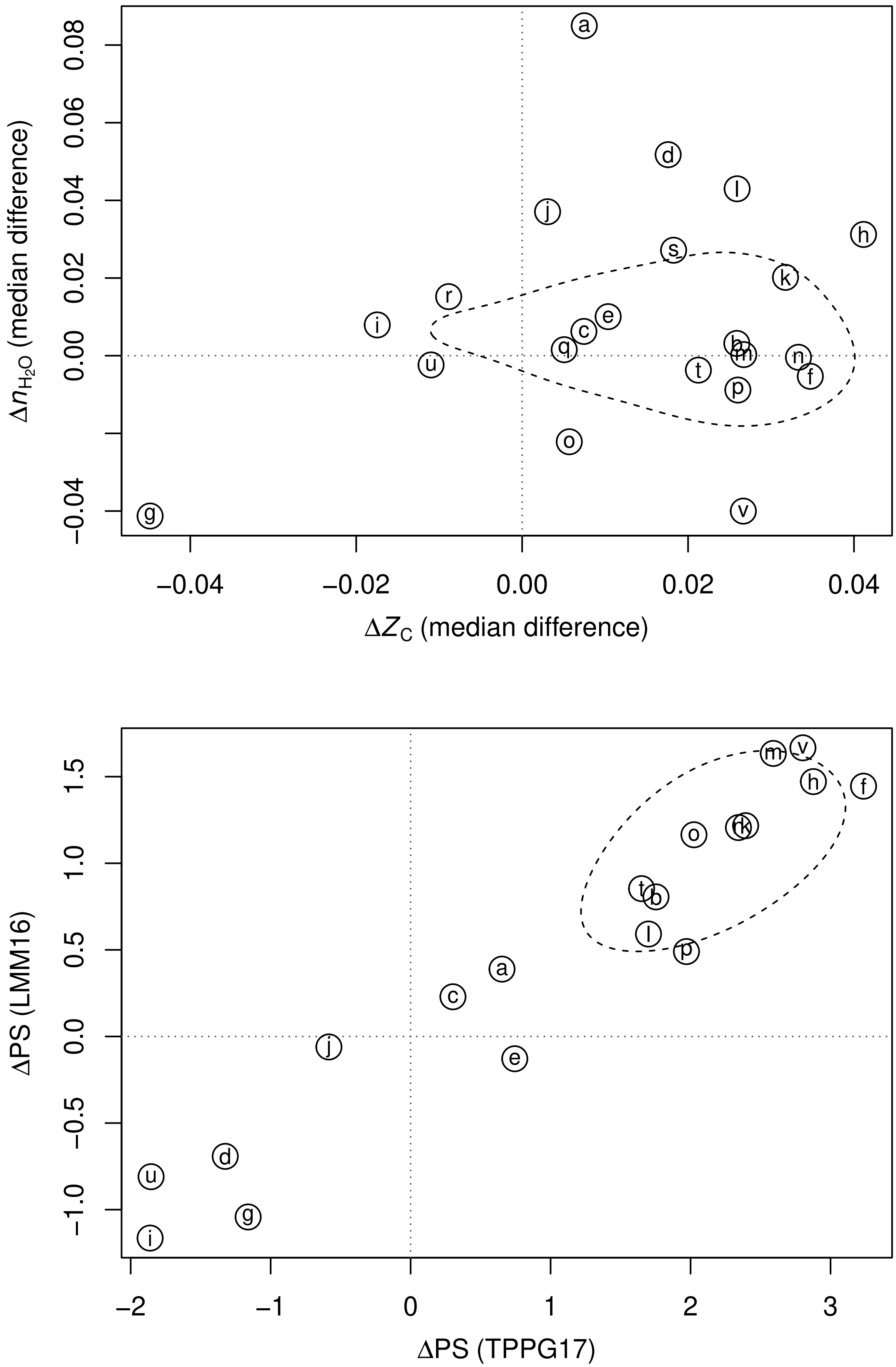


**Fig. S15.** Compositional analysis and phylostrata for differentially expressed proteins in pancreatic cancer.

### Legend for Fig. S15

Abbreviations: T – tumor; N – normal; LCM – laser capture microdisecction; FFPE – formalin-fixed paraffin-embedded; DM – diabetes mellitus.

| Set | Reference | Description | Down | Up |
|---|---|---|---|---|
| a | LHE+04 | T / adjacent N | 41 | 69 |
| b | CYD+05 | T / N | 60 | 88 |
| c | CGB+05 | T / N | 48 | 54 |
| d | CTZ+09 | T / adjacent N | 28 | 29 |
| e | MLC+11 | T / adjacent N | 38 | 45 |
| f | PCS+11 | FFPE T / N | 207 | 152 |
| g | TMW+11 | accessible T / N | 108 | 86 |
| h | KBK+12 | FFPE T / N | 38 | 47 |
| i | KHO+13 | T / N | 78 | 57 |
| j | KPC+13 | T / adjacent N | 257 | 456 |
| k | WLL+13a | T / adjacent N with DM | 208 | 219 |
| l | WLL+13a | T / adjacent N without DM | 56 | 167 |
| m | YKK+13 | T / adjacent N | 84 | 83 |
| n | ZNWL13 | LCM T / adjacent N | 227 | 148 |
| o | ISI+14 | T / adjacent N | 65 | 34 |
| p | BZQ+14 | T / matched N | 55 | 98 |
| q | MZH+14 | mouse tumor / healthy | 38 | 28 |
| r | BHB+15 | mouse organoids T / N | 486 | 526 |
| s | KKC+16 | mouse 10 w T / N | 37 | 108 |
| t | CHO+18 | T / adjacent N | 109 | 129 |
| u | SWW+18 | T / adjacent N | 284 | 324 |
| v | ZAH+19 | T / N | 89 | 76 |

**a**. Tables 2 and 3 of (178). **b**. Tables 1 and 2 of (179). **c**. Table 2 of (180). **d**. Table 1 of (181). **e**. IPI numbers from Supplementary Table S2 of (182). **f**. Supplementary Table 3 of (183). **g**. Extracted from the SI Table of (184). **h**. Supplementary Tables 2 and 3 of (185). **i**. SI Table S3 of (186), filtered to include proteins with an expression ratio >2 [or <0.5] in at least 5 of the 7 experiments and ratio >1 [or <1] in all experiments. **j**. Supplementary Table 2 of (187). **k**. **l**. Supplementary Tables S3 and S4 of (188), including proteins with >3/2 or <2/3 fold change in at least 3 of 4 iTRAQ experiments for different pooled samples. **m**. Supplementary Tables 2 and 3 of (189) (data file provided by Youngsoo Kim). **n**. SI Table S5 of (190). **o**. SI Table S5 of (191), filtered to exclude proteins marked as "not passed", i.e. having inconsistent regulation. **p**. Table S6, Sheet 2 of (192). **q**. Table 1 of (193). **r**. Table S6 of (194). **s**. Supplementary Table of (195). **t**. Supplementary Table S3 of (196). **u**. Table S1 of (197), filtered to exclude proteins with opposite expression changes in different patients. **v**. Gene names extracted from Figure 1b of (198).

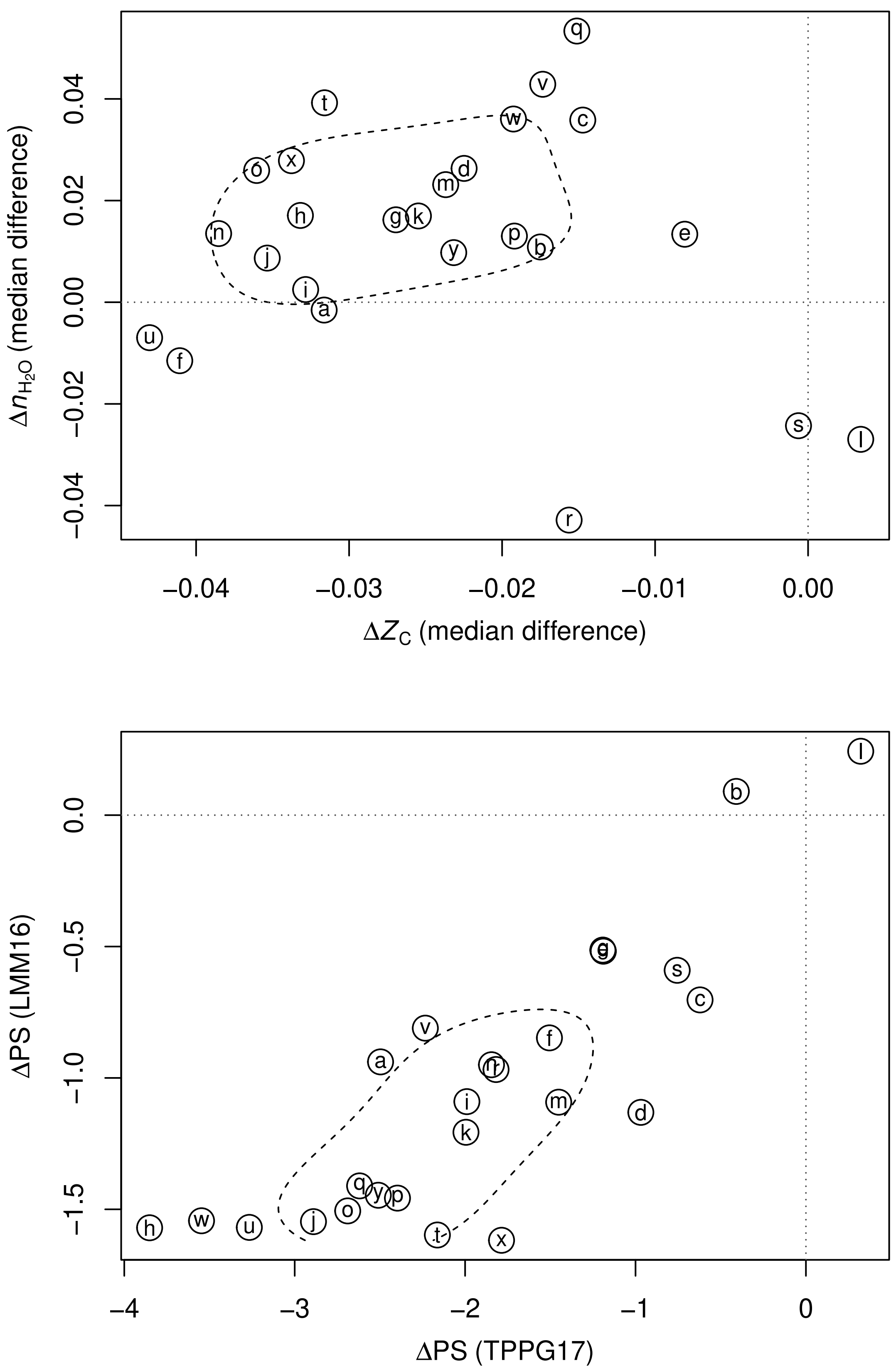


**Fig. S16.** Compositional analysis and phylostrata for differentially expressed proteins in prostate cancer.

### Legend for Fig. S16

Abbreviations: PCa – prostate cancer; BPH – benign prostatic hyperplasia; FFPE – formalin-fixed paraffin-embedded; OCT – optimal cutting temperature compound-embedded; TMA – tissue microarray; GS – Gleason Score; CiRT – common internal retention time.

| Set | Reference | Description | Down | Up |
|---|---|---|---|---|
| a | GTR+08 | PCa / BPH | 35 | 29 |
| b | KPB+10 | PCa / adjacent benign | 68 | 69 |
| c | HZH+12 | PCa / adjacent benign Protein | 22 | 37 |
| d | JHZ+13 | PCa / adjacent benign | 23 | 36 |
| e | LCS+14 | glycoproteins PCa / normal | 82 | 138 |
| f | CZL+16 | protein expression bioinformatics | 74 | 157 |
| g | IWT+16 | FFPE tumor / adjacent benign | 223 | 426 |
| h | GLZ+18 | acinar PCa / matched BPH | 87 | 114 |
| i | GLZ+18 | ductal PCa / matched BPH | 347 | 303 |
| j | LAJ+18 | PC / BPH | 273 | 220 |
| k | LAJ+18 | CRPC / BPH | 313 | 193 |
| l | MAN+18 | PCa / BPH | 44 | 33 |
| m | KRN+19 | PCa G1 / BPH | 32 | 43 |
| n | KRN+19 | PCa G2 / BPH | 30 | 22 |
| o | KRN+19 | PCa G3 / BPH | 43 | 24 |
| p | KRN+19 | PCa G4 / BPH | 44 | 51 |
| q | KRN+19 | PCa G5 / BPH | 22 | 24 |
| r | MMF+19 | FFPE PCa / adjacent benign GS=6 | 40 | 136 |
| s | MMF+19 | FFPE PCa / adjacent benign GS=6 and GS>=8 | 37 | 143 |
| t | TOT+19 | FFPE TMA PCa / normal | 36 | 86 |
| u | ZYW+19 | OCT LG PCa / adjacent normal | 143 | 54 |
| v | ZYW+19 | OCT HG PCa / adjacent normal | 215 | 94 |
| w | KHN+20 | PCa / control | 227 | 125 |
| x | SHC+20 | FFPE PCa / BPH | 58 | 308 |
| y | ZZX+20 | tumor / normal CiRT | 104 | 1451 |

**a**. Table 2 of (199). **b**. Supplementary Tables 3 and 4 of (200). **c**. Table S2 of (201). **d**. Table S1 of (202). **e**. Supplemental Table S7 of (203). **f**. Supplementary Table 1 (204) (proteins recorded with only Down-regulation or Up-regulation). **g**. Supplementary Table S3 of (205), filtered to include proteins listed with FDR < 0.1. **h**. **i**. Extracted from Table S4 (206). Values were quantile normalized, then ratios were calculated between the median values for each cancer type (acinar and ductal) and corresponding normal tissue; ratios > 1.5 or < 2/3 were used to identify differentially expressed proteins. **j**. **k**. Extracted from Supplementary Data 1 of (207) (sheet “Area-proteins”) by applying quantile normalization to peak areas then calculating median values across all runs and samples for each of BPH , PC (primary prostate cancer), and CRPC (castration resistant prostate cancer). A cutoff of 2-fold in ratios of medians (PC / BPH or CRPC / BPH) was used to identify differentially expressed proteins. **l**. Table 2 of (208). **m**. **n**. **o**. **p**. **q**. Supporting Information Table S3 of (209) (G1–G5: PCa grades). **r**. **s**. Tables S4d (GS = 6) and S4f (GS ≥ 8 or GS = 6) of (210) (GS: Gleason Score). **t**. Supplementary Table S3a of (211). **u**. **v**. Table S5 of (212) (LG: low-grade PCa; HG: high-grade PCa). **w**. Supplemental Table S2 of (213), filtered to include proteins with $\log_2$ fold change > 1 or < -1 in at least one experiment. **x**. Table S2, Sheet D:S3-SWATH_protein_matrix of (214), filtered to include proteins quantified in at least 50% of both tumor and normal samples and with median fold change > 2 or < 0.5. **y**. Supplementary Tables S3E–S3F of (215).

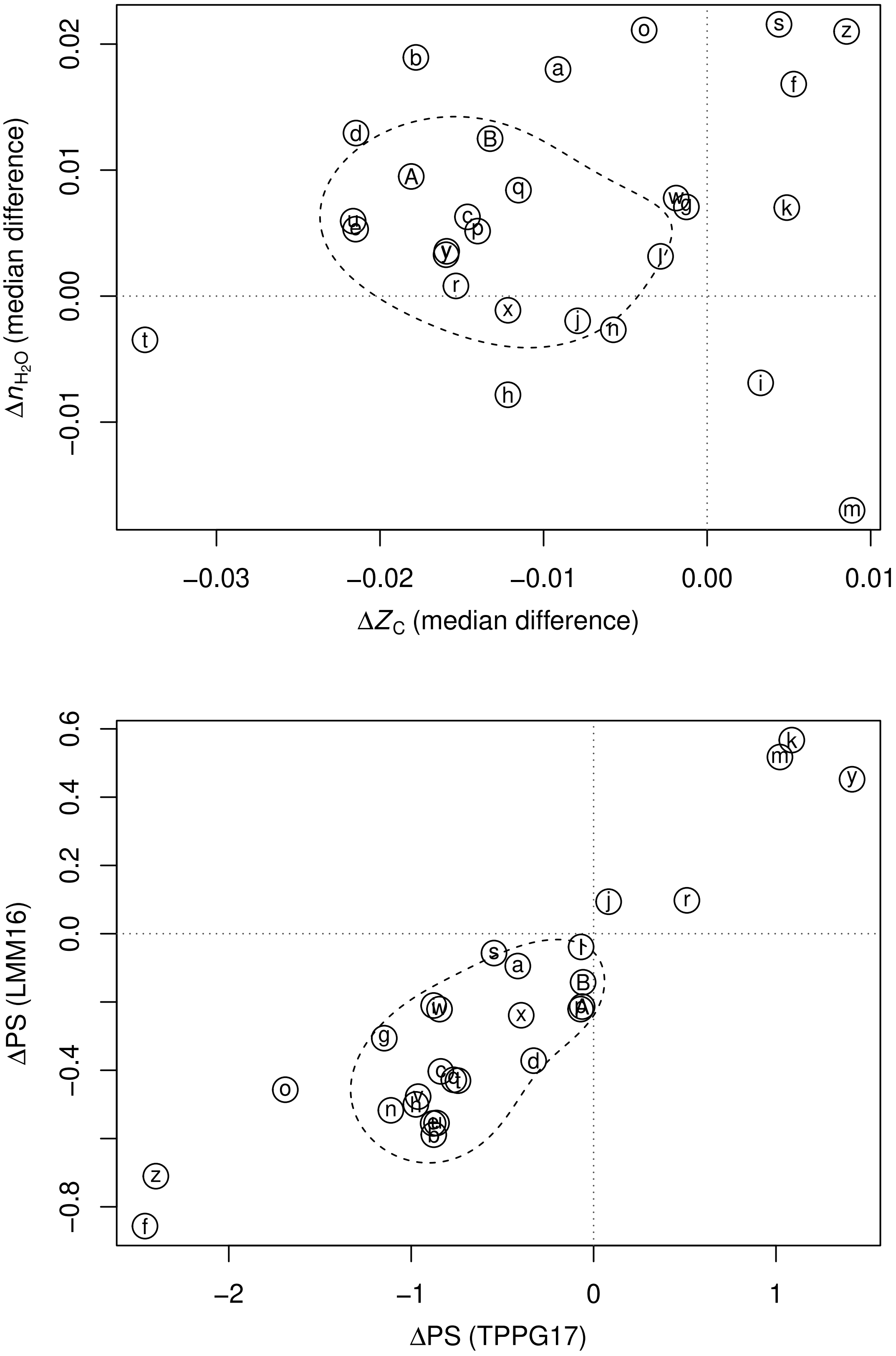


**Fig. S17.** Compositional analysis and phylostrata for proteins corresponding to differentially expressed genes between TCGA and GTEx datasets.

### Legend for Fig. S17

| Set | Reference | Description | Down | Up |
|---|---|---|---|---|
| a | GEPIA2 | ACC | 2013 | 391 |
| b | GEPIA2 | BLCA | 1722 | 735 |
| c | GEPIA2 | BRCA | 1749 | 1256 |
| d | GEPIA2 | CESC | 2972 | 1627 |
| e | GEPIA2 | COAD | 2165 | 2327 |
| f | GEPIA2 | DLBC | 771 | 7887 |
| g | GEPIA2 | ESCA | 1088 | 2401 |
| h | GEPIA2 | GBM | 1883 | 4642 |
| i | GEPIA2 | HNSC | 557 | 1385 |
| j | GEPIA2 | KICH | 3002 | 615 |
| k | GEPIA2 | KIRC | 1141 | 1433 |
| l | GEPIA2 | KIRP | 1258 | 827 |
| m | GEPIA2 | LAML | 2578 | 2490 |
| n | GEPIA2 | LGG | 1376 | 3500 |
| o | GEPIA2 | LIHC | 570 | 1342 |
| p | GEPIA2 | LUAD | 2430 | 949 |
| q | GEPIA2 | LUSC | 3167 | 1654 |
| r | GEPIA2 | OV | 3567 | 2241 |
| s | GEPIA2 | PAAD | 335 | 8167 |
| t | GEPIA2 | PRAD | 1778 | 518 |
| u | GEPIA2 | READ | 2331 | 2455 |
| v | GEPIA2 | SKCM | 3007 | 2252 |
| w | GEPIA2 | STAD | 708 | 3458 |
| x | GEPIA2 | TGCT | 6227 | 2174 |
| y | GEPIA2 | THCA | 2341 | 524 |
| z | GEPIA2 | THYM | 783 | 10099 |
| A | GEPIA2 | UCEC | 3834 | 1733 |
| B | GEPIA2 | UCS | 3418 | 1592 |

**a**. – **B**. GEPIA2 (216).

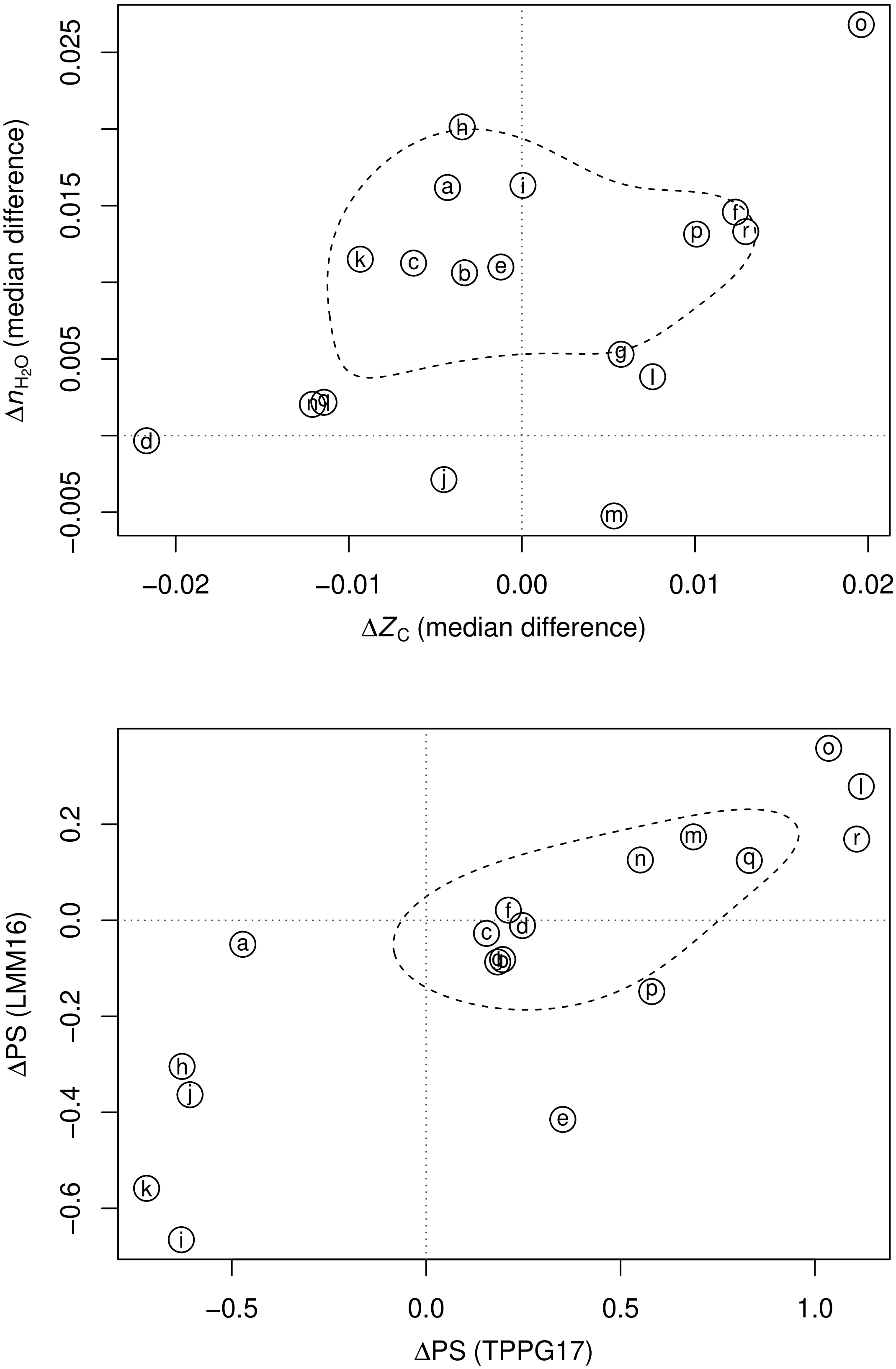


**Fig. S18.** Compositional analysis and phylostrata for differentially expressed proteins in HPA datasets.

**Legend for Fig. S18**

| Set | Reference | Description | Down | Up |
|---|---|---|---|---|
| a | HPA19 | breast cancer / breast | 77 | 512 |
| b | HPA19 | cervical cancer / cervix, uterine | 367 | 207 |
| c | HPA19 | colorectal cancer / colon | 263 | 423 |
| d | HPA19 | endometrial cancer / endometrium 1 | 117 | 263 |
| e | HPA19 | glioma / cerebral cortex | 95 | 209 |
| f | HPA19 | head and neck cancer / salivary gland | 1075 | 534 |
| g | HPA19 | liver cancer / liver | 51 | 423 |
| h | HPA19 | lung cancer / lung | 402 | 268 |
| i | HPA19 | lymphoma / lymph node | 446 | 69 |
| j | HPA19 | melanoma / skin 1 | 180 | 516 |
| k | HPA19 | ovarian cancer / ovary | 151 | 1075 |
| l | HPA19 | pancreatic cancer / pancreas | 361 | 293 |
| m | HPA19 | prostate cancer / prostate | 621 | 197 |
| n | HPA19 | renal cancer / kidney | 939 | 97 |
| o | HPA19 | stomach cancer / stomach 1 | 1963 | 111 |
| p | HPA19 | testis cancer / testis | 1202 | 123 |
| q | HPA19 | thyroid cancer / thyroid gland | 784 | 387 |
| r | HPA19 | urothelial cancer / urinary bladder | 1050 | 94 |

**a**. – **r**. Human Protein Atlas (217).

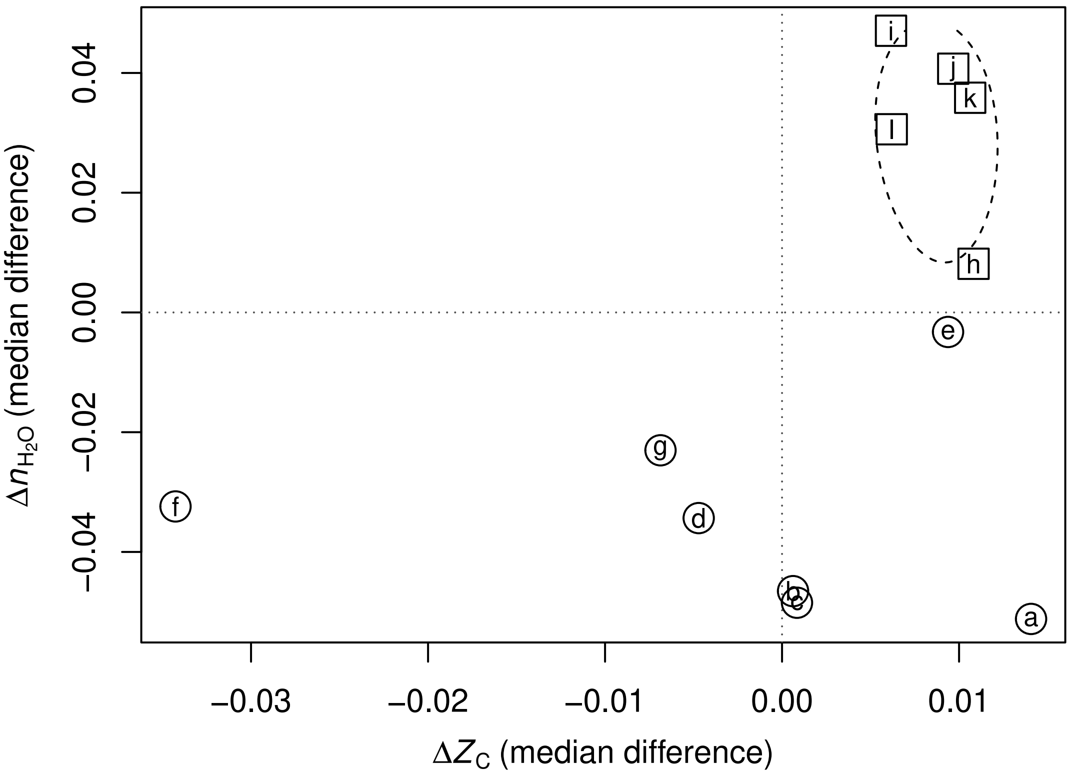


**Fig. S19.** Compositional analysis for proteins corresponding to differentially expressed genes in hypoosmotic and hyperosmotic shock in *Saccharomyces cerevisiae*. Circles and squares represent hyperosmotic and hypoosmotic conditions, respectively.

**Legend for Fig. S19**

| Set | Reference | Description | Down | Up |
|---|---|---|---|---|
| a | GSK+00 | yeast transcriptome in 1M sorbitol at 5 min | 141 | 212 |
| b | GSK+00 | yeast transcriptome in 1M sorbitol at 15 min | 802 | 923 |
| c | GSK+00 | yeast transcriptome in 1M sorbitol at 30 min | 430 | 613 |
| d | GSK+00 | yeast transcriptome in 1M sorbitol at 45 min | 84 | 286 |
| e | GSK+00 | yeast transcriptome in 1M sorbitol at 60 min | 30 | 98 |
| f | GSK+00 | yeast transcriptome in 1M sorbitol at 90 min | 73 | 153 |
| g | GSK+00 | yeast transcriptome in 1M sorbitol at 120 min | 119 | 220 |
| h | GSK+00 | yeast transcriptome in hypoosmotic shock at 5 min | 292 | 332 |
| i | GSK+00 | yeast transcriptome in hypoosmotic shock at 15 min | 479 | 459 |
| j | GSK+00 | yeast transcriptome in hypoosmotic shock at 30 min | 718 | 386 |
| k | GSK+00 | yeast transcriptome in hypoosmotic shock at 45 min | 502 | 327 |
| l | GSK+00 | yeast transcriptome in hypoosmotic shock at 60 min | 312 | 258 |

**a**. – **l**. complete_dataset.txt of (218), filtered to include genes with an expression ratio > $\log_2(1.5)$ or < $\log_2(1/1.5)$ at any time in the "1M sorbitol" or "Hypo-osmotic shock" experiments.

reveals drastic changes in fatty acid metabolism and plasma membrane transporters. *Journal of Proteome Research* 14(9):4005–4018.

ulated protein complexes in primary prostate cancer. *Clinical Proteomics* 16(1):15.
213. Kwon OK, et al. (2020) Identification of novel prognosis and prediction markers in advanced prostate cancer tissues based on quantitative proteomics. *Cancer Genomics - Proteomics* 17(2):195–208.
214. Sun R, et al. (2020) Accelerated protein biomarker discovery from FFPE tissue samples using single-shot, short gradient microflow SWATH MS. *Journal of Proteome Research* in press.
215. Zhu T, et al. (2020) DPHL: A pan-human protein mass spectrometry library for robust biomarker discovery. *bioRxiv* preprint.
216. Tang Z, Kang B, Li C, Chen T, Zhang Z (2019) GEPIA2: An enhanced web server for large-scale expression profiling and interactive analysis. *Nucleic Acids Research* 47(W1):W556–W560.
217. Uhlén M, et al. (2015) Tissue-based map of the human proteome. *Science* 347(6220):1260419.
218. Gasch AP, et al. (2000) Genomic expression programs in the response of yeast cells to environmental changes. *Molecular Biology of the Cell* 11(12):4241–4257.
